# Synthetic glabridin derivatives mitigate steatohepatitis in a diet-induced biopsy-confirmed non-alcoholic steatosis hepatitis mouse model through paraoxonase-2

**DOI:** 10.1101/2021.10.01.462722

**Authors:** Gu-Choul Shin, Hyeong Min Lee, Na Yeon Kim, Sang-Ku Yoo, Yun Sun Park, Hyung Soon Park, Dongryeol Ryu, Kwang Pyo Kim, Kyun-Hwan Kim

## Abstract

Limited therapeutic agents have been developed for non-alcoholic steatohepatitis (NASH), a common immunometabolic disease that can progress to hepatic cirrhosis and cancer. Glabridin and its derivatives are potential therapeutics for some metabolic diseases. However, the therapeutic effects of glabridin and its derivatives on NASH and their biological functions are unclear. This study demonstrated the role of synthetic glabridin derivatives (SGDs) in alleviating hepatic steatosis and inflammation in a biopsy-confirmed rodent NASH model. SGDs exerted therapeutic effects by activating autophagy and the antioxidant defense system, which mitigate NASH pathogenesis. The cellular target of HSG4112, an SGD, was paraoxonase 2. These findings will enable the development of novel therapeutics for NASH in the future.

## Introduction

Non-alcoholic steatohepatitis (NASH), which is characterized by hepatic inflammation and fibrosis, can progress to hepatic cirrhosis or cancer.^1^ The hepatic accumulation of free fatty acids (FFAs) and toxic lipid intermediates promotes the development of non-alcoholic fatty liver disease (NAFLD) by inducing oxidative stress, mitochondrial dysfunction, and hepatic inflammation.^2–4^ These metabolic disorders along with the consumption of high-calorie diet and sedentary lifestyle are a major public health concern worldwide.^5,6^ Thus, there is an urgent need to develop therapeutic agents for NASH that can effectively promote toxic lipid metabolite catabolism and mitigate oxidative stress and hepatic inflammation.

Clinical trials have examined various therapeutic drugs for NASH. Semaglutide, a glucagon-like peptide-1 receptor agonist (GLP1-RA), exhibited promising results for the clinical treatment of NASH in a recent phase 2 trial. However, semaglutide was not effective against the fibrosis stage.^7^ The mechanism underlying the pharmacological activity of semaglutide involves the suppression of gastric emptying and food intake through GLP-1R-dependent regulation of the physiology in both the hypothalamus and hindbrain. Semaglutide treatment decreases bodyweight and liver weight, alleviates hepatic steatosis, and inhibits the onset of NAFLD in mouse models.^8^ However, hepatocytes do not express the canonical GLP-1R,^9^ and the expression of GLP-1R has not been conclusively established in other liver cells. The role of GLP-1RA in the attenuation of NASH is unclear. The elucidation of a novel mechanism distinct from that of GLP-1RA may provide novel insights into the alleviation of NASH.

Glabridin has important applications in the food, dietary supplement, and cosmetic industries. However, the application of glabridin is limited owing to its low water solubility, low bioavailability, and unpredictable stability.^10^ Orally administered glabridin exhibits various bioactivities, including antioxidant, anti-inflammatory, and estrogen-like activities, and promotes lipid catabolism.^11,12^ Thus, glabridin and its derivatives are potential therapeutic agents for immunometabolic diseases, such as NASH although the underlying mechanisms have not been completely elucidated.

The chemical stability and oral bioavailability of synthetic glabridin derivatives (SGDs), which were recently synthesized by our group, are higher than those of glabridin. In this study, the therapeutic effect of SGDs on NASH was evaluated using a rodent model. Additionally, the underlying mechanism of action (MOA) of SGDs was elucidated.

## Results

### HSG4112 alleviates steatohepatitis in the biopsy-confirmed NASH mouse model

The therapeutic efficacy of SGDs in progressive steatohepatitis was examined using an amylin diet-induced obesity (AMLN-DIO) mouse model, which exhibits severe inflammation and liver fibrosis. The liver of mice fed on AMLN diet for 37 weeks was biopsied 4 weeks before SGD treatment to confirm the development of NASH. Mice were administered with SGDs during the last 6 weeks of AMLN diet supplementation (**Supplementary Fig. 1a**). Hematoxylin and eosin (H&E) staining and oil red O staining revealed that steatosis in the HSG4112-treated and HSG4113-treated groups was significantly alleviated when compared with that in the vehicle-treated group (**Fig. 1a**). HSG4112 and HSG4113 mitigated the AMLN diet-induced liver fibrosis (Sirius red staining and Lgals3 and Acta2 immunostaining) and intrahepatic monocyte infiltration (Adgre1 immunostaining). Consistently, biochemical analysis revealed that HSG4112 and HSG4113 mitigated the AMLN diet-induced upregulation of serum and hepatic cholesterol levels and plasma levels of Fgf21 and Lep (steatosis markers), Tnfa, and Il10 (inflammatory markers), and alanine aminotransferase (ALT), aspartate aminotransferase (AST), and alkaline phosphatase (ALP) (hepatic injury markers) but did not affect the Il5, Il6, and Cxcl1 levels (**Fig. 1b** and **Supplementary Fig. 2a**).

**Fig. 1.**
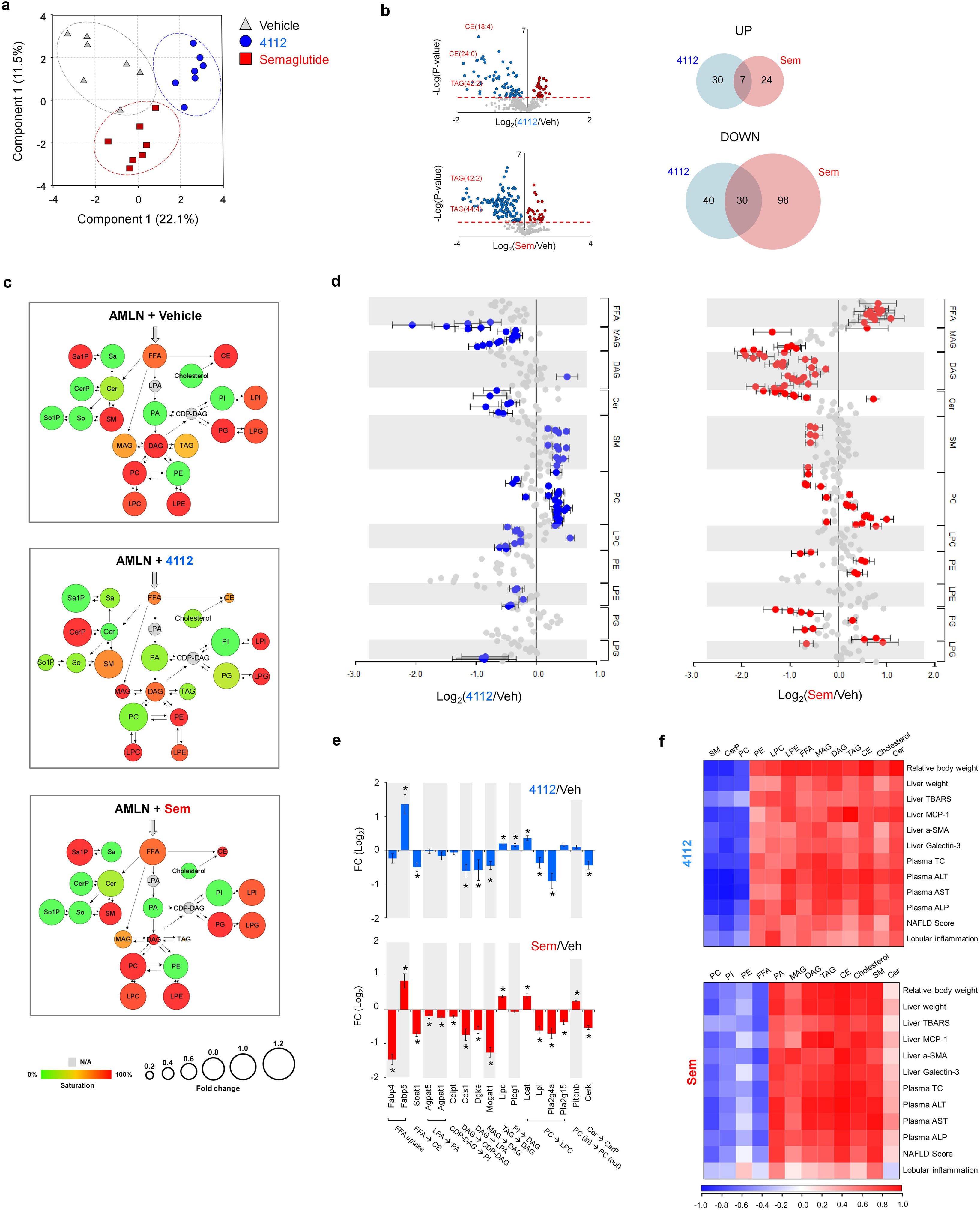
Synthetic glabridin derivatives (SGDs) mitigate steatosis and inflammation in a biopsy-confirmed rodent non-alcoholic steatosis hepatitis (NASH) model. **a** Mice were fed with amylin (AMLN) diet and NASH was confirmed using biopsy. Next, the mice were orally administered with HSG4112 (100 mg/kg bodyweight) and HSG4113 (200 mg/kg bodyweight) for 6 weeks once daily. Representative results of hematoxylin and eosin, oil red O, Sirius red staining and Lgals3, Acta, Adgre1 immunostaining (scale bar = 50 μm). The histopathological scores are shown above. **b** Quantification of plasma total cholesterol, liver cholesterol, and serum Tnfa, alanine aminotransferase, aspartate aminotransferase, and alkaline phosphatase. **c–f** Histological scores of non-alcoholic fatty liver disease activity (c), lobular inflammation (d), steatosis (e), and ballooning degeneration (f) in biopsies before treatment (pre) and after treatment (post) are shown in the left. Numbers of animals exhibiting exacerbation (higher), no change (same), and alleviation (lower) of disease after treatment relative to before treatment are shown in the right. **g–i** Effect of SGDs on bodyweight (g), liver weight (h), and cumulative food intake (i). Data are presented as mean ± standard deviation (n = 12 for the vehicle-treated group; n = 11 for the HSG4112-treated group; n = 10 for the HSG4113-treated group). In **a, b**, and **h**, data of the SGD-treated and vehicle-treated groups were compared using one-way analysis of variance, following by Dunnett’s post-hoc test. In **c–f**, data were analyzed using the Fisher’s test, followed by the Bonferroni test.

The NASH-related phenotypes before SGD treatment were compared with thoseat week 6 post-SGD treatment. Compared with that at the baseline, the NAFLD activity score (NAS), which indicates the degree of lobular inflammation, steatosis, and ballooning degeneration, was lower at week 6 post-HSG4112 treatment. In contrast, the NAS of the HSG4113-treated group was not significantly different at baseline and week 6 post-treatment (**Fig. 1c–e**, **Supplementary Fig. 3**). Compared with that in the vehicle-treated group, the number of mice exhibiting decreased NAS was significantly higher in the HSG4112-treated group and similar in the HSG4113-treated group. HSG4112 and HSG4113 did not alleviate hepatic ballooning and fibrosis (**Fig. 1f** and **Supplementary Fig. 2b**). Additionally, HSG4112 significantly decreased the liver weight and bodyweight (**Fig. 1g–h**). The HSG4112-treated, HSG4113-treated, and vehicle groups exhibited similar daily food intake (**Fig. 1i**). These findings indicate that HSG4112 alleviated NASH.

### HSG4112 alleviates hepatic inflammation and NASH without decreasing the appetite

A phase 2 clinical trial revealed that semaglutide alleviates NASH.^13^ Thus, the therapeutic effects of HSG4112 and semaglutide on NASH were comparatively analyzed (**Supplementary Fig. 1b**). Similar to the 6-week treatment regimen, long-term (10 weeks) treatment with HSG4112 alleviated hepatic steatosis in the AMLN-DIO mouse model (**Fig. 2a**) and downregulated the cholesterol levels (**Fig. 2b**) to levels observed in the semaglutide-treated group. Semaglutide and HSG4112 mitigated liver fibrosis and intrahepatic monocyte infiltration (**Fig. 2a**) and downregulated Ccl2 (an inflammatory marker) and ALT, AST, and ALP (hepatic injury markers) (**Fig. 2b**).

**Fig. 2.**
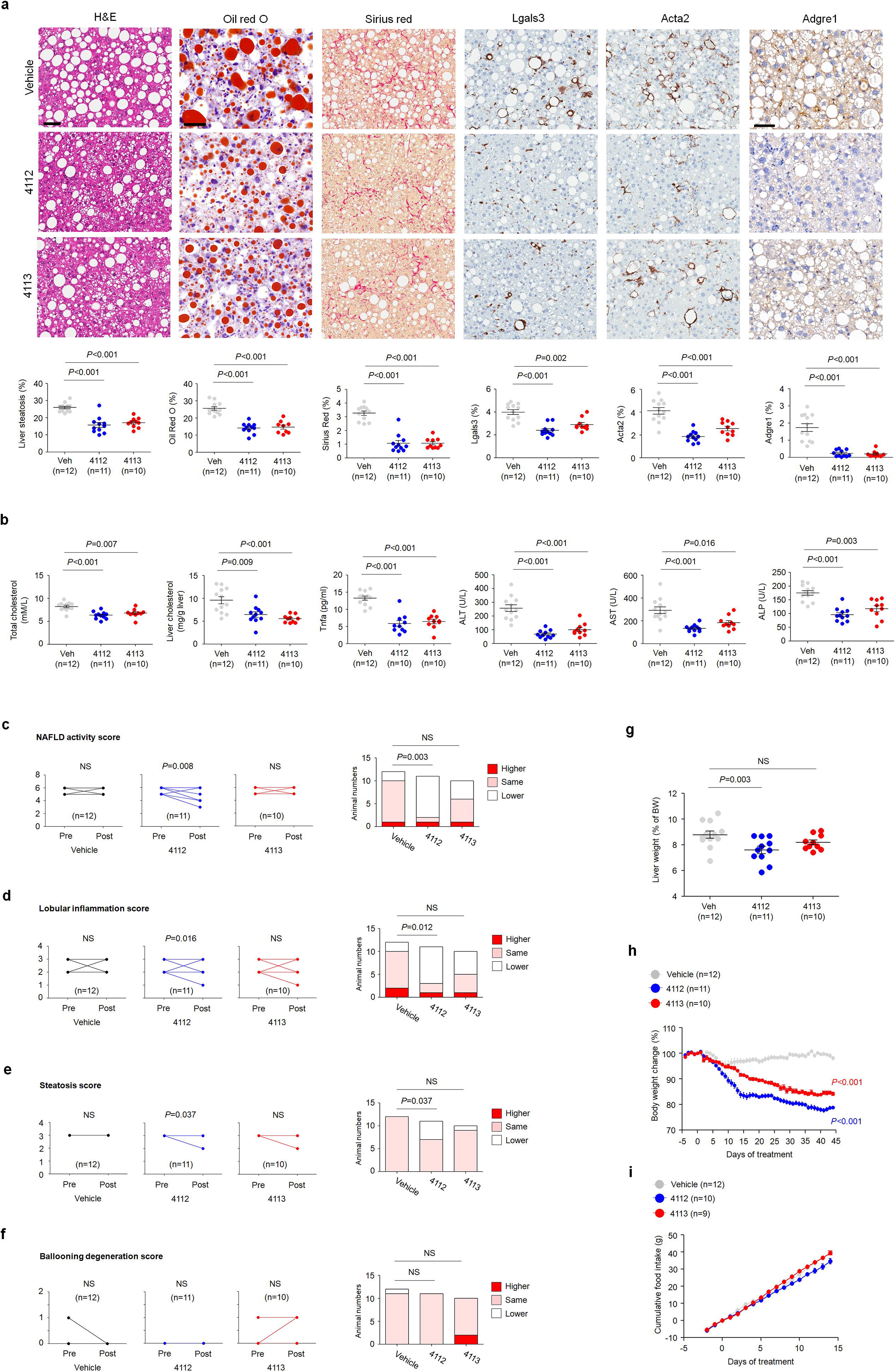
Therapeutic effect of HSG4112 on non-alcoholic steatosis hepatitis (NASH) is similar to that of semaglutide. **a** Amylin (AMLN) diet-fed mice were treated once daily with HSG4112 (50 or 100 mg/kg bodyweight, oral administration) and semaglutide (30 nM/kg bodyweight, subcutaneous injection) for 10 weeks. Representative results of hematoxylin, oil red O, and Sirius red staining, Lgals3, Acta2, and Adgre1 immunostaining (scale bars = 50 μm), and histopathological scores are shown above. **b** Quantification of plasma total cholesterol, liver cholesterol, liver Ccl2, and serum alanine aminotransferase (ALT), aspartate aminotransferase (AST), and alkaline phosphatase (ALP). **c** and **d** Histological scores of NAFLD activity (c) and lobular inflammation (d) in biopsies before treatment (pre) and after treatment (post) are shown on the left. Numbers of animals exhibiting exacerbation (higher), no change (same), and alleviation (lower) of disease after treatment relative to before treatment are shown right. **e–i** Effect of HSG4112 and semaglutide on the liver weight (e), bodyweight (f), fat/non-fat mass (g), muscle weight (h), and cumulative food intake (i). Data are presented as mean ± standard deviation (n = 15 for the vehicle-treated group; n=15 for the HSG4112 (50 mg/kg bodyweight)-treated group; n = 14 for the HSG4112 (100 mg/kg bodyweight)-treated group; n = 14 for the semaglutide-treated group). In **a, b, e, g, h**, and **I,** data of the HSG4112-treated group vs. vehicle-treated group and semaglutide-treated group vs. vehicle-treated were compared using one-way analysis of variance, followed by Dunnett’s post-hoc test. In **c** and **d**, data were compared using the Fisher’s test, followed by the Bonferroni test.

The mice were stratified into HSG4112-treated, semaglutide-treated, and vehicle groups based on the NASH phenotypes at the baseline. The NASH phenotypes at the baseline were compared with those at week 10 post-drug treatment. Treatment with HSG4112 and semaglutide for 10 weeks significantly decreased the NAS (**Fig. 2c**). Additionally, the number of mice with decreased NAS in the HSG4112-treated and semaglutide-treated groups was significantly higher than that in the vehicle-treated group. In contrast to semaglutide, HSG4112 mitigated lobular inflammation (**Fig. 2d**). However, HSG4112 and semaglutide did not mitigate hepatic fibrosis (**Supplementary Fig. 4**). HSG4112 and semaglutide significantly decreased the liver weight and bodyweight after treatment for 10 weeks (**Fig. 2e–f** and **Supplementary Fig. 5a**). The initial bodyweight reduction rate in the semaglutide-treated group was higher than that in the HSG4112-treated group. However, the final bodyweight was similar in both groups (**Fig. 2f**). Additionally, HSG4112 and semaglutide alleviated the AMLN diet-induced changes in body composition and adiposity (**Fig. 2g–h** and **Supplementary Fig. 5b–c**). In contrast to HSG4112, semaglutide markedly decreased the daily food intake (**Fig. 2i** and **Supplementary Fig. 5d**). These findings indicate that HSG4112 alleviates NASH without decreasing the appetite and mitigates hepatic inflammation.

### Semaglutide and HSG4112 differentially affected the hepatic lipid profiles

Next, the mechanisms underlying the therapeutic effects of HSG4112 and semaglutide were examined. The hepatic lipid profiles of the HSG4112-treated and semaglutide-treated groups were examined using liquid chromatography-electrospray ionization-tandem mass spectrometry (LC-ESI-MS/MS). Principal component analysis (PCA) revealed that the lipid profiles of the HSG4112-treated, semaglutide-treated, and vehicle-treated groups exhibited distinct clustering. This indicated that the MOA of HSG4112 was distinct from that of semaglutide (**Fig. 3a**). Volcano plot analysis revealed that HSG4112 downregulated 70 lipid species, including cholesterol esters (CEs, 24:0 and 18:4) and triacylglycerols (TAGs, 42:2), while semaglutide downregulated 128 lipid species, mainly TAGs (42:2 and 44:4) (**Fig. 3b**). HSG4112 significantly upregulated 37 lipid species, including sphingomyelins (SMs) and phosphatidylcholines (PCs), while semaglutide upregulated 31 lipid species, including PCs and FFAs. The relative contents and saturation levels of various glycerolipids, including monoacylglycerol (MAG), diacylglycerol (DAG), and TAG, were markedly downregulated in the HSG4112-treated and semaglutide-treated groups (**Fig. 3c** and **Supplementary Fig. 6**). Additionally, HSG4112 downregulated FFAs, ceramide (Cer), and lysophospholipid (LPL) species, including lyso-phosphatidylcholine (LPC), lyso-phosphatidylethanolamine (LPE), lyso-phosphatidylinositol (LPI), and lyso-phosphatidylglycerol (LPG), whereas semaglutide downregulated phosphatidic acid.

**Fig. 3.**
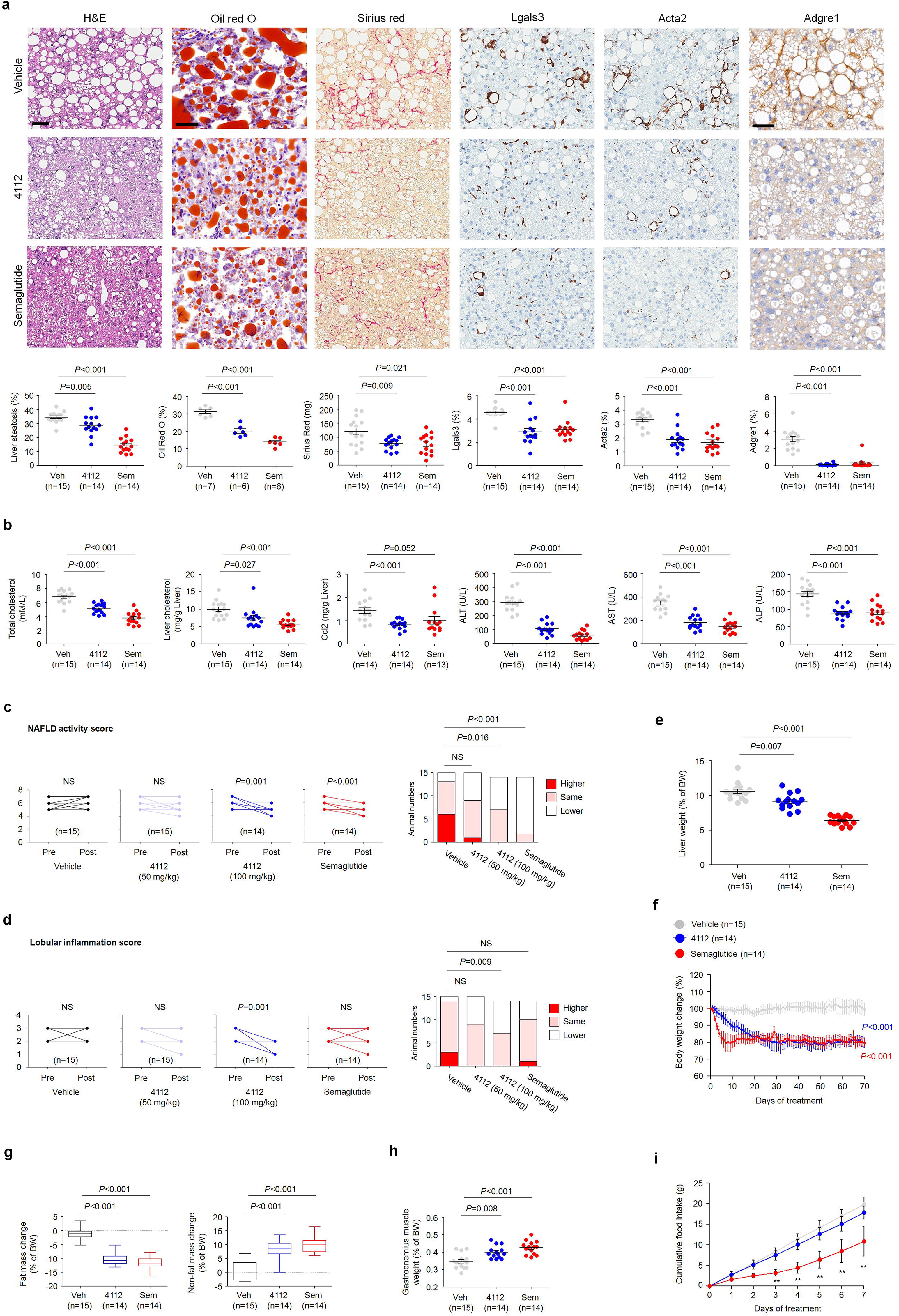
HSG4112 and semaglutide differentially affect the lipid profiles. **a** Principal coordinate analysis of hepatic levels of total lipids in the vehicle-treated, HSG4112-treated, and semaglutide-treated amylin (AMLN) diet-fed mice. **b** Volcano plot analysis of the hepatic levels of total lipids in the HSG4112-treated (left panel) and semaglutide-treated (right panel) groups when compared with those in the vehicle-treated group. Venn diagram showing the number of differentially regulated hepatic lipids in the HSG4112-treated and semaglutide-treated group relative to the vehicle-treated group. **c** Analysis of hepatic lipid classes in the vehicle-treated, HSG4112-treated, and semaglutide-treated groups. In the liver of the vehicle-treated group (top panel), the schematic diagram shows the relative levels of incorporation of exogenous fatty acids into sphingolipids and glycerophospholipids. Lipid classes identified using liquid chromatography-tandem mass spectrometry analysis are presented as color-coded circles. The lipid classes were designated as saturated if all their fatty acid chains were saturated and as unsaturated if they had at least one unsaturated fatty acid chain. The percentage of saturated lipid species is shown for each class from green (low saturation) to red (high saturation). Lipid classes not identified are shown in gray. The size of the circles is set to the arbitrary unit of 1 for the control cells. The hepatic lipid species in the HSG4112-treated and semaglutide-treated groups are presented in the middle and bottom panels, respectively. The size of the circle is proportional to the fold change in the content of lipid species in the drug-treated group relative to the vehicle-treated group. **d** Forest plots showing individual hepatic lipid species in the HSG4112-treated and semaglutide-treated AMLN diet-fed mice expressed as a fold change relative to the vehicle-treated group. Plots in blue or red represent differential individual lipid species between the drug-treated and vehicle-treated groups; p < 0.05. **e** Analysis of the levels of enzymes involved in hepatic lipid metabolism in the HSG4112-treated and semaglutide-treated AMLN diet-fed mice when compared with those in the vehicle-treated group. **f** Heatmap analysis of the correlation between lipid classes and pathological features in the HSG4112-treated and semaglutide-treated AMLN diet-fed mice. Each cell is color-coded and the Pearson’s correlation coefficient is shown. The legend for color-coding is shown below. LPAs, lyso-phosphatidic acids; PAs, phosphatidic acids; MAG, monoacylglycerol; DAG, diacylglycerol; TAG, triacylglycerol; PC, phosphatidylcholine; LPC, lyso-phosphatidylcholine; PE, phosphatidylethanolamine; LPE, lyso-phosphatidylethanolamine; PI, phosphatidylinositol; LPI, lyso-phosphatidylinositol; PG, phosphatidylglycerol; LPG, lyso-phosphatidylglycerol; Cer, ceramide; SM, sphingomyelin; CerP, ceramide-1-phosphate; Sa, sphinganine; Sa1P, sphinganine-1-phosphate; So, sphingosine; So1P, sphingosine-1-phosphates. Fabp4, fatty acid-binding protein 4; Fabp5, fatty acid-binding protein 5; Soat1, sterol O-acyltransferase 1; Agpat5, 1-acylglycerol-3-phosphate O-acyltransferase 5; Agpat1, 1-acylglycerol-3-phosphate O-acyltransferase 1; Cdipt, CDP-diacylglycerol-inositol 3-phosphatidyltransferase; Cds1, CDP-diacylglycerol synthase 1; Dgke, diacylglycerol kinase epsilon; Mogat1, monoacylglycerol O-acyltransferase 1; Lipc, lipase C hepatic type; Plcg1, phospholipase C gamma 1; Lcat, lecithin-cholesterol acyltransferase; Lpl, lipoprotein lipase; Pla2g4a, phospholipase A2 group IVA; Pla2g15, phospholipase A2 group XV; Pitpnb, phosphatidylinositol transfer protein beta; CerK1, ceramide kinase 1. *p < 0.05. Error bars indicate standard deviation.

The levels of 78 individual lipid species were significantly different between the HSG4112-treated and semaglutide-treated groups (**Fig. 3d** and **Supplementary Fig. 7**). HSG4112 downregulated most individual MAGs, whereas semaglutide downregulated most individual DAGs and TAGs. In contrast to semaglutide, HSG4112 downregulated most individual LPCs, LPEs, and LPGs and upregulated individual SMs. HSG4112 distinctively downregulated some FFAs, including long-chain FFAs, whereas semaglutide upregulated most FFAs, including saturated and unsaturated FFAs. The levels of cholesterol species and PCs were not markedly different between the HSG4112-treated and semaglutide-treated groups but were significantly different from those in the vehicle-treated group. Lipidome analysis revealed that the MOA of HSG4112 was distinct from that of semaglutide.

Transcriptomic analysis was performed to examine the expression levels of lipid catabolic enzymes. The mRNA levels of genes encoding enzymes involved in FFA uptake, CE biosynthesis from FFAs, cytidine diphosphate (CDP)-DAG biosynthesis from DAG, lysophosphatidic acid (LPA) biosynthesis from DAG, DAG biosynthesis from MAG and TAG, and ceramide-1-phosphate (CerP) biosynthesis from Cer were not significantly different between the HSG4112-treated and semaglutide-treated groups (**Fig. 3e**). However, the levels of *Agpat5* and *Agpat1*, which encode enzymes involved in phosphatidic acid biosynthesis from LPA, were significantly downregulated in the semaglutide-treated group but not in the HSG4112-treated group. The levels of *Lpl* and *Pla2g15*, which are involved in LPC biosynthesis from PC, were significantly downregulated in the semaglutide-treated group. Furthermore, the expression of *Pitpnb*, which is involved in the release of extracellular PC, was significantly upregulated in the semaglutide-treated group but not in the HSG4112-treated group.

Two clusters of lipid species were strongly correlated with alleviating NASH in the HSG4112-treated group. The levels of these lipid species in the HSG4112-treated and semaglutide-treated groups were comparatively analyzed. A cluster of 10 lipid species, including PE, LPC, LPE, FFA, and Cer, was positively correlated with NASH alleviation in the HSG4112-treated group. In the semaglutide-treated group, 8 lipid species containing phosphatidic acid and SM were positively correlated with NASH alleviation (**Fig. 3f**). The LPC, LPE, FFA, and Cer levels were positively correlated with the alleviation of hepatic inflammation in the HSG4112-treated group but not in the semaglutide-treated group. These results suggest that the regulatory effects of HSG4112 on lipid catabolism are distinct from those of semaglutide.

### HSG4112 modulates the autophagic and anti-inflammatory pathways

To investigate the mechanisms underlying HSG4112-mediated NASH alleviation, the gene expression levels were analyzed using RNA sequencing (RNAseq). Of the 22028 genes, the levels of approximately 10% of genes in the semaglutide-treated (2244 genes) and HSG4112-treated (2413 genes) groups were different from those in the vehicle group (**Fig. 4a**). To identify the specific target genes of the drugs, the differentially expressed genes (DEGs) between the HSG4112-treated and semaglutide-treated groups were analyzed. Among the meta-signature genes, HSG4112 and semaglutide co-regulated 1146 genes, which represent common gene signatures of the drugs for NASH alleviation (**Fig. 4b**). Furthermore, HSG4112 and semaglutide specifically altered the expression of 1267 and 1098 genes, respectively, which represent the drug-specific gene signatures. Gene Ontology (GO) analysis revealed that 384 upregulated overlapping genes were significantly enriched in biological oxidation, metabolism of lipids, steroid biosynthesis, and fatty acid degradation, whereas 762 downregulated overlapping genes were significantly enriched in extracellular matrix organization, immune system, liver fibrosis, and inflammation signaling (**Fig. 4c–d** and **Supplementary Fig. 8a–d**). Meanwhile, 656 genes specifically upregulated in the HSG4112-treated group were related to glutathione metabolism, branched-chain amino acid degradation, and autophagy, while 491 genes specifically upregulated in the semaglutide-treated group were related to bile acid and arachidonic acid metabolism (**Fig. 4c and e**). Furthermore, 611 genes specifically downregulated in the HSG4112-treated group were related to cell adhesion molecules, antigen processing and presentation, and MAPK signaling pathway, whereas 607 genes specifically downregulated in the semaglutide-treated group were related to Rho GTPase cycle and fibrin clot formation (**Fig. 4c and f**).

**Fig. 4.**
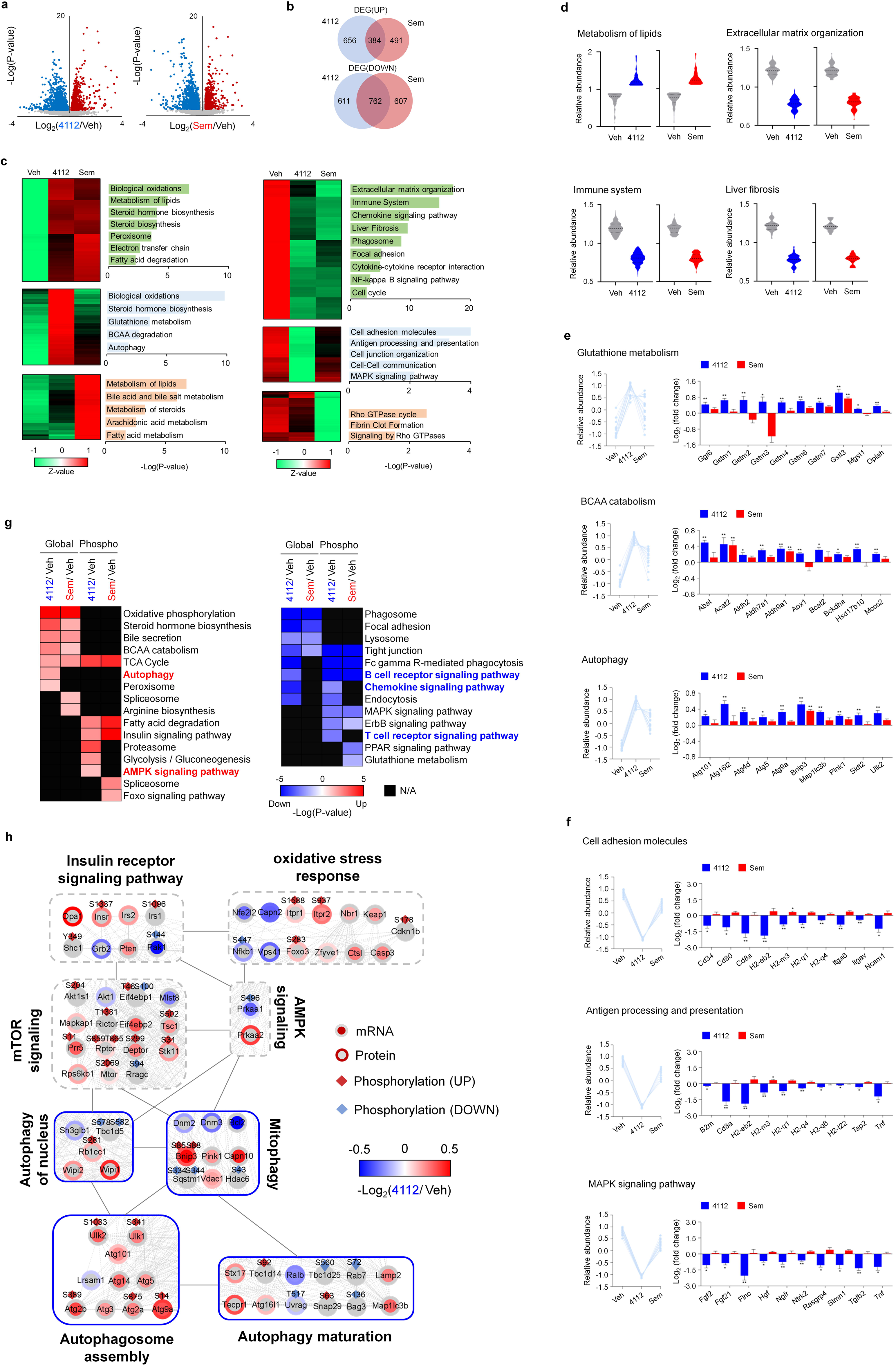
Transcriptome and phosphoproteome analyses indicated that HSG4112 exerts therapeutic effects by promoting autophagy activation. **a** Volcano plot of hepatic differentially expressed genes (DEGs) between the HSG4112-treated/semaglutide-treated and vehicle-treated amylin (AMLN) diet-fed mice. Genes upregulated or downregulated by more than 2-fold are shown in red and blue, respectively. **b** Venn diagram showing the number of hepatic DEGs between the HSG4112-treated/semaglutide-treated and vehicle-treated groups. **c** Two-dimensional hierarchical clustering of DEGs between different pairs of HSG4112-treated, semaglutide-treated, and vehicle-treated groups. The figure shows the most significant Gene Ontology (GO) biological processes for each cluster of genes upregulated (left panel) and downregulated (right panel) in the HSG4112-treated and semaglutide-treated groups (individually or in both groups) relative to the vehicle-treated group. Red and green denote highly and weakly expressed genes, respectively. **d** Violin plots showing the mean and variance of overlapping gene clusters between drug-treated and vehicle-treated groups. **e** and **f** Among HSG4112-specific gene clusters, the relative abundance scores of upregulated (e) and downregulated (f) genes were compared between all pairs of HSG4112-treated, semaglutide-treated, and vehicle-treated groups. Bar plots showing the average fold change in expression of perturbed genes in HSG4112-specific gene clusters. **g** Top-ranked pathways in phosphoproteome analysis that are significantly altered in the HSG4112-treated and semaglutide-treated groups relative to the vehicle-treated group. Red cells represent the pathways enriched by proteins exhibiting upregulated phosphorylation (Phospho) and corresponding protein expression (Global) in the drug-treated group relative to the vehicle-treated group. Blue cells represent the pathways enriched by the proteins exhibiting downregulated phosphorylation and corresponding total protein expression in the drug-treated groups relative to the vehicle-treated group. Each cell is color-coded based on the fold change in the expression of genes in the drug-treated groups relative to the vehicle-treated group. The legend for the color codes is shown below. Bold words represent major signaling pathways enriched by HSG4112-specific phosphoproteins. **h** Network scheme showing the interactions of genes involved in the autophagy signaling pathway. The genes/proteins are selected from the DEGs of the transcriptome and phosphoproteome analysis with enhanced connectivity and significant differential expression in the HSG4112-treated group. Each gene/protein symbol is color-coded based on the fold change in expression of genes/proteins in the drug-treated groups relative to the vehicle-treated groups. The legend for color-coding is shown below and represented with the phosphorylation site. *p < 0.05. Error bars indicate standard deviation. N/A, not available.

Next, the levels of phosphoproteins and their corresponding total proteins in the HSG4112-treated and semaglutide-treated groups were comparatively analyzed. Common signatures of upregulated phosphoproteins and their corresponding total proteins were associated with oxidative phosphorylation, steroid hormone biosynthesis, and fatty acid degradation (**Fig. 4g**). Meanwhile, common signatures of downregulated phosphoproteins and their corresponding total proteins were associated with the phagosome, focal adhesion, and tight junction. Compared with that in the semaglutide-treated group, the phosphorylation of autophagy-linked AMPK signaling pathway-related components was upregulated, while that of B cell receptor, chemokine, and T cell receptor signaling pathway-related components was downregulated in the HSG4112-treated group.

To examine the network of biological processes related to specific gene and protein signatures in the HSG4112-treated group, functional interaction networks were constructed (**Fig. 4h**). The autophagy-linked biological processes were the major targets of HSG4112. Insulin receptor signaling, oxidative stress response, mTOR signaling, and AMPK signaling are the upstream signals of autophagy. HSG4112 activated autophagy initiation signaling (Ulk1 and Ulk2), autophagosome biogenesis (Rb1cc1, Wipi1, Wipi2, and Atgs), and autophagy maturation (Map1Lc3b, Stx17, Tecpr1, and Lamp2). These findings suggest that HSG4112 regulates lipid catabolism through the induction of the autophagy pathways.

### HSG4112 activates autophagy/mitophagy and attenuates mitochondrial oxidative stress under lipotoxic conditions

Next, the effect of HSG4112 on alleviating lipotoxicity in hepatocytes was examined. HSG4112 significantly mitigated FFA-induced lipid accumulation in the hepatocytes (**Fig. 5a**). Omics data indicated that HSG4112 promoted autophagy, which promotes the degradation of lipid droplets and the release of FFAs. Lipid droplets and FFAs are then oxidized through mitochondrial fatty acid oxidation (FAO), which results in the alleviation of lipotoxicity.^14^ Hence, the autophagic flux was examined by monitoring the conversion of LC3B-I to LC3B-II and the degradation of Sqstm1. Consistent with the results of the mechanistic study on hepatic lipid profiles (**Fig. 3c and e**), compared with those in the FFA-treated cells, the LC3B levels were upregulated and the Sqstm1 levels were downregulated in the HSG4112-treated cells (**Fig. 5b**). HSG4112-induced autophagy activation was confirmed using the autophagy indicator tandem fluorescence-tagged LC3B. Compared with that in the control and FFA-treated groups, the number of cells with autophagic flux (red fluorescent protein (RFP)) was higher in the HSG4112-treated group (**Fig. 5c**). HSG4112-induced autophagic flux was confirmed using the lysosomal fusion inhibitor bafilomycin A (BFA). The number of autophagosomes in the BFA-treated cells was markedly higher than that in the control or FFA-treated cells as evidenced by the accumulation of both green fluorescent protein (GFP) and RFP puncta (**Fig. 5d**). Map1lc3b and Becn1 immunostaining analyses revealed that HSG4112 significantly promoted autophagy activation in the mouse liver (**Fig. 5e**).

**Fig. 5.**
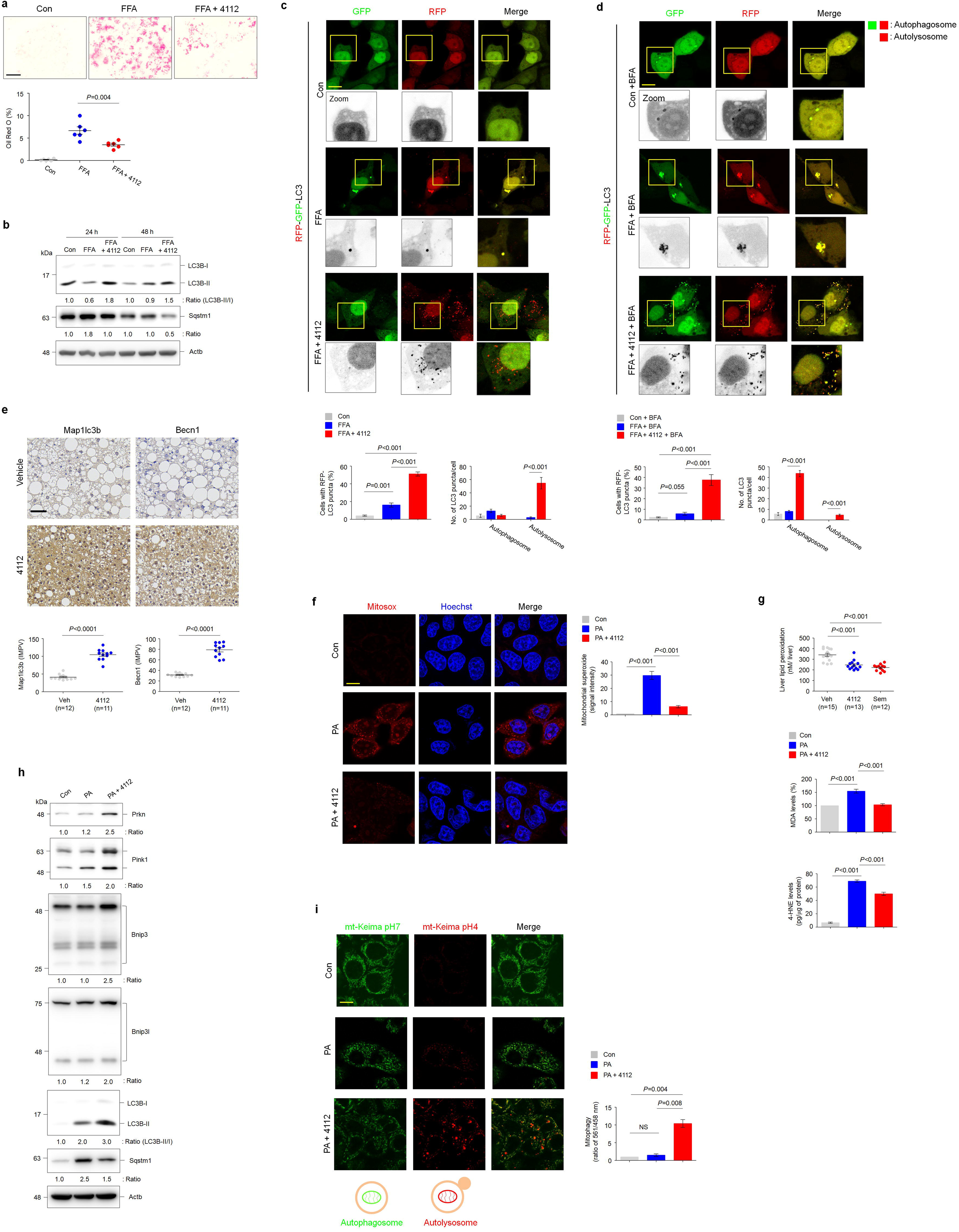
HSG4112 activates autophagy/mitophagy and alleviates lipotoxic oxidative stress. **a** L02 hepatocytes were treated with free fatty acids (FFAs) (500 μM) alone or in combination with HSG4112 (3 μM) 72 h and stained with oil red O to determine lipid accumulation. Representative images from six independent experiments are shown above (scale bar = 50 μm). Quantification of oil red O staining intensities is shown below. **b** Immunoblotting analysis of autophagy flux activation-related markers (Map1lc3b, Sqstm1, and Actb) in L02 cells treated with FFAs alone or in combination with HSG4112 for indicated durations. Actb was used as a loading control. Band intensities of indicated proteins are shown below. **c** Confocal fluorescence analysis showing the puncta of tandem fluorescent probe-tagged LC3B (red fluorescent protein (mRFP)-green fluorescent protein (GFP)-LC3B). L02 cells were transfected with mRFP-GFP-LC3B plasmid and treated with FFAs alone or in combination with HSG4112 for 72 h. Cells with both red and green fluorescent puncta (autophagosome) and those with only red fluorescent puncta (autolysosome) were quantified. Representative images from three independent experiments are shown above (scale bar = 20 μm). Bar plots of the average cell numbers with red fluorescent puncta (autophagy) or average numbers of autophagosome or autolysosome per cell are shown below. Error bars indicate standard deviation. **d** Confocal fluorescence analysis showing the puncta of mRFP-GFP-LC3B in cells treated with bafilomycin A (BFA), an autophagy maturation inhibitor. L02 cells transfected with mRFP-GFP-LC3B plasmid were treated with FFAs alone or in combination with HSG4112 for 72 h, followed by treatment with BFA (100 nM) for 5 h. Representative images from three independent experiments are shown above (scale bar = 20 μm). Bar plots of the average cell numbers exhibiting autophagy or average numbers of autophagosome or autolysosome per cell are shown below. Error bars indicate standard deviation. **e** Immunohistochemical staining analysis of hepatic Map1lc3b and Becn1 levels in HSG4112-treated amylin (AMLN) diet-fed mice. Representative images of hepatic sections of the vehicle-treated (n = 12) and HSG4112-treated (n = 11) mice are shown above (scale bar = 50 μm). Quantification of staining intensity (inverted mean pixel values) is shown below. **f** Confocal fluorescence images showing the generation of mitochondrial superoxide, which was analyzed using MitoSox staining. L02 cells were treated with palmitic acid (PA) (125 μM) alone or in combination with HSG4112 (3 μM) for 24 h and stained with MitoSox. Cell nuclei were stained with Hoechst 33342. Representative images from three independent experiments are shown in the left (scale bar = 20 μm). Bar plots of the quantification of MitoSox staining intensities are shown right. Error bars indicate standard deviation. **g** Quantification of lipid peroxidation in the liver and hepatocytes. Hepatic lipid peroxidation in the HSG4112-treated (n = 13), semaglutide-treated (n = 12), and vehicle-treated (n = 15) AMLN diet-fed mice was measured using the malondialdehyde (MDA) accumulation assay (top panel). Bar plots of the average lipid peroxidation in hepatocytes determined using the MDA (middle panel) and 4-hydroxynonenal assays (bottom panel). Lipid peroxidation in L02 cells treated with PA alone or in combination with HSG4112 for 24 h. Data were obtained from three independent experiments. Error bars indicate standard deviation. **h** Immunoblotting analysis of mitophagy activation-related markers (Prkn, Pink1, Bnip3, Bnip3l, Map1lc3b, Sqstm1, and Actb) in L02 cells treated with PA alone or in combination with HSG4112 for 5 h. Actb was used as a loading control. Band intensities of indicated proteins are shown below. **i** Confocal fluorescence analysis of mitophagy activation-related markers in hepatocytes. L02 cells were transfected with mt-Keima, a pH-dependent fluorescent mitophagy probe, were treated with PA alone or in combination with HSG4112 for 24 h and the mt-Keima fluorescent intensity was quantified. Green fluorescent filaments (pH 7.0) of mt-Keima indicate mitochondrial networks, while enlarged red fluorescence of mt-Keima (pH 4.0) indicates mitophagy flux (fused with lysosome). Representative images from three independent experiments are shown in the left (scale bar = 20 μm). Bar plots of the average red (excitation 561 nm) and green (excitation 458 nm) fluorescence intensities are shown on the right. Error bars indicate standard deviation. NS, non-significant.

The analysis of genes involved in electron transport chain revealed that HSG4112 upregulated mitochondrial FAO (**Supplementary Fig. 8f**). FAO is reported to enhance reactive oxygen species (ROS) generation.^15^ Hence, the effect of HSG4112 on mitochondrial oxidative stress generated during FAO was examined. HSG4112 markedly downregulated palmitic acid (PA)-induced mitochondrial superoxide to the level observed in the control group (**Fig. 5f**). Additionally, HSG4112 significantly downregulated lipid peroxidation in the NASH mouse liver and PA-treated hepatocytes (**Fig. 5g**). Mitophagy is a quality control mechanism that protects the cells against cytotoxic ROS production by clearing damaged mitochondria.^16^ HSG4112 suppressed the PA-induced mitophagy inhibition by upregulating Prkn/Pink1, Bnip3, and Bnip3l. Additionally, HSG4112 promoted the mitophagic flux by upregulating LC3B-II and downregulating Sqstm1 (**Fig. 5h**). The mitophagy rates in hepatocytes stably expressing mt-Kemia, a mitochondrial dual-excitation ratiometric fluorescent protein reporter of mitophagy, were analyzed.^59^ Consistent with results of immunoblotting analysis, mitophagy activation in the HSG4112-treated group was higher than that in the PA-treated or control groups as evidenced by enhanced mitochondrial localization to acidic lysosome (**Fig. 5i**).^17^ These findings indicate that HSG4112 activates the autophagy-FAO axis and attenuates oxidative stress by activating mitophagy.

### HSG4112 activates autophagy and alleviates oxidative stress through Pon2

To elucidate the mechanism of HSG4112, the intracellular localization of HSG4112 was examined using Alexa 488 azide-conjugated propargylated HSG4112, which was synthesized using the Click-iT reaction. Fluorescence analysis revealed that HSG4112 primarily localized to the mitochondria (**Fig. 6a**). Hepatic gene function analysis indicated that HSG4112 targets mitochondrial functions (**Supplementary Fig. 8e**). To identify the target protein of HSG4112, biotin-labeled HSG4112 was synthesized using the Click-iT reaction. Cell lysates were subjected to the pull-down assay and chemical proteome analysis. The enriched GO terms for 130 HSG4112-bound proteins in the mitochondria were related to the regulation of overall mitochondrial energy metabolism, including oxidative phosphorylation, FAO, tricarboxylic acid cycle, and oxidative stress homeostasis (**Supplementary Fig. 9a–b**). HSG4112 interacted with paraoxonase 2 (Pon2), which is reported to hydrolyze lipid peroxide and alleviate oxidative stress (**Fig. 6b**).^18^ To confirm the specific interaction, a competitive pull-down assay was performed by pre-incubating the cell lysates with free HSG4112. Free HSG4112 dose-dependently decreased the interaction between HSG4112 and Pon2. This was consistent with the results of studies using purified recombinant Pon2 (rPon2) protein (**Fig. 6c**). The interaction between HSG4112 and Pon2 may modulate the Pon2 esterase and lactonase activities. Hence, the Pon2 activity in HSG4112-treated cells was examined. HSG4112 mitigated the PA-induced downregulation of Pon2 activity although it did not affect the Pon2 expression (**Fig. 6d**). Next, the ability of HSG4112 to specifically activate Pon2 was examined as Pon1 and Pon3 also exhibit esterase and lactonase activities in hepatocytes. HSG4112 did not affect the esterase and lactonase activities in *Pon2* knockdown (KD) cells, which suggested that HSG4112 specifically activates Pon2 (**Fig. 6e**). Additionally, HSG4112 mitigated the oxidized linoleic acid-induced downregulation of rPon2 activity *in vitro* (**Fig. 6f**). These results indicate that HSG4112 suppresses lipid peroxide-induced inhibition of Pon2 activity. Furthermore, Pon2 is reported to exhibit antioxidant activities through the reduction of superoxide generated by CoQ10 in the mitochondria, which leads to the downregulation of lipid peroxidation.^19–21^ Thus, the effect of HSG4112 on mitochondrial superoxide was examined. HSG4112 markedly downregulated mitochondrial superoxide production in wild-type cells but not in *Pon2* KD cells (**Fig. 6g**). Consistently, HSG4112 did not downregulate lipid peroxidation in *Pon2* KD cells (**Fig. 6h**), which indicated that Pon2 activity is crucial for the antioxidant effect of HSG4112. The intrinsic function of Pon2 in autophagy/mitophagy activation is unclear. However, the potential role of Pon2 in HSG4112-mediated activation of autophagy/mitophagy was examined in this study. Pon2 KD inhibited autophagy/mitophagy under basal conditions (**Fig. 6i–j**) as evidenced by Sqstm1 upregulation and Prkn and Bnip3l downregulation. The effect of HSG4112 on FFA-induced or PA-induced autophagy/mitophagy was abolished in *Pon2* KD cells. These results indicate that Pon2 contributes to the HSG4112-mediated activation of autophagy. HSG4112 did not decrease lipid accumulation in *Pon2* KD cells (**Fig. 6k**). These findings suggest that HSG4112 activates autophagy/mitophagy and alleviates lipotoxicity in hepatocytes through Pon2.

**Fig. 6.**
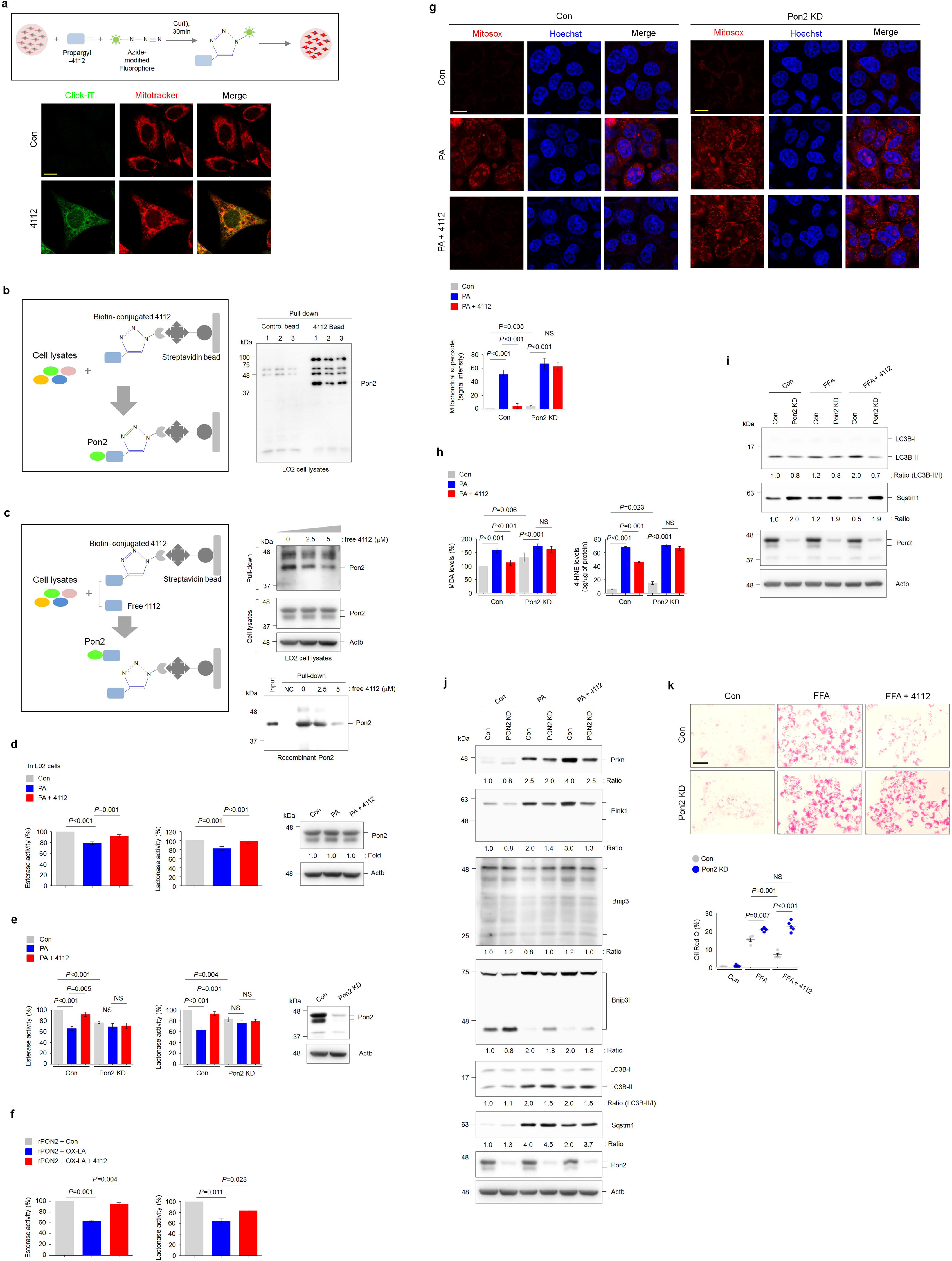
HSG4112 activates mitophagy and autophagy, reduces lipid accumulation, and maintains redox homeostasis through Pon2. **a** Confocal fluorescence images showing the mitochondrial localization of HSG4112. L02 cells were treated with Alexa488-azide-conjugated propargyl-HSG4112 (3 μM), which was prepared using the Click-iT reaction buffer kit, for 5 h. Mitochondria were stained with MitoTracker Red. The experimental procedure is depicted above. Representative images from three independent experiments are shown below (scale bar = 20 μm). **b** Identification of target proteins of HSG4112 using chemical proteome analysis. Proteins extracted from L02 cells interacted with the propargyl-HSG4112-biotin-streptavidin bead complex. The experimental procedure is depicted on the left. Immunoblotting analysis of the interaction between HSG4112 and Pon2 using three independent samples is shown on the right. **c** Competitive binding assay to determine the specific interaction between HSG4112 and Pon2. Cell lysates or purified recombinant Pon2 protein were incubated with different concentrations of free HSG4112, followed by incubation with streptavidin bead-conjugated HSG4112. The experimental procedure is depicted on the left. Immunoblotting analysis of the specific interaction between HSG4112 and Pon2 is shown on the right. **d** Results of the enzymatic assay demonstrating the esterase and lactonase activities of Pon2 in hepatocytes. Pon2 activity in L02 cells treated with palmitic acid (PA) alone or in combination with HSG4112 for 24 h. Bar plots of the average esterase and lactonase activities are shown on the left. Data were obtained from three independent experiments. Error bars indicate standard deviation. Immunoblot analysis of Pon2 and Actb is shown on the right. Actb was used as a loading control. Band intensities of the indicated proteins are shown below. **e** Pon2 activities in the *Pon2* knockdown (KD) and control L02 cells treated with PA alone or in combination with HSG4112 for 24 h. Bar plots of the average esterase and lactonase activities are shown on the left. Data were obtained from three independent experiments. Error bars indicate standard deviation. Immunoblotting analysis of Pon2 and Actb is shown on the right. Actb was used as a loading control. NS, non-significant. **f** Purified recombinant Pon2 activities incubated with oxidized linoleic acid (OX-LA) alone or in combination with HSG4112 for 10 min. Bar plots of the average esterase and lactonase activities of recombinant Pon2. Data were obtained from three independent experiments. Error bars indicate standard deviation. **g** Confocal fluorescence images showing the generation of mitochondrial superoxide, which was analyzed using MitoSox. *Pon2* KD and control L02 cells were treated with PA alone or in combination with HSG4112 for 24 h and stained with MitoSox. Cell nuclei were stained with Hoechst 33342. Representative images from three independent experiments are shown above (scale bar = 20 μm). Bar plots of the MitoSox staining intensity are shown below. Error bars indicate standard deviation. **h** The quantification of lipid peroxidation in *Pon2* KD and control L02 cells treated with PA alone or in combination with HSG4112 for 24 h. Bar plots of the average lipid peroxidation determined using the malonaldehyde accumulation (left panel) and 4-hydroxynonenal assays (right panel). Data were obtained from three independent experiments. Error bars indicate standard deviation. NS, non-significant. **i** Immunoblotting analysis of autophagy flux activation-related markers (Map1lc3b, Sqstm1, and Actb) in *Pon2* KD and control L02 cells treated with free fatty acids (FFAs) alone or in combination with HSG4112 for 48 h. Actb was used as a loading control. The band intensities of indicated proteins are shown below. **j** Immunoblotting analysis of mitophagy activation-related markers (Prkn, Pink1, Bnip3, Bnip3l, and Map1lc3b) in *Pon2* KD and control L02 cells treated with PA alone or in combination with HSG4112 for 24 h. Actb was used as a loading control. Band intensities of indicated proteins are shown below. **k** *Pon2* KD and control L02 cells were treated with FFAs alone or in combination with HSG4112 combined for 72 h and stained with oil red O to examine lipid accumulation. Representative images from five independent experiments are shown above (scale bar = 50 μm). Quantification of oil red O staining intensity is shown below. NS, non-significant.

## Discussion

Glabridin is reported to be a potential therapeutic for metabolic diseases, such as obesity and diabetes mellitus.^22,23^ However, the effect of glabridin and SGDs on NASH severity has not been examined. In this study, a clinically relevant biopsy-confirmed NASH mouse model was used to accurately examine the therapeutic effect of SGDs on NASH.^24,25^ The anti-inflammatory effects of HSG4112 were higher than those of semaglutide. Moreover, HSG4112 mitigated the AMLN diet-induced adverse effects on the overall physical attributes, including bodyweight loss, liver weight loss, and enhanced lean mass, without decreasing the appetite. HSG4112 markedly reprogrammed the hepatic transcription and metabolism associated with lipid catabolism, hormone biosynthesis, protein anabolism, redox homeostasis, and immune response. GLP-1RAs, such as semaglutide may indirectly alleviate NASH as the canonical GLP-1R is not expressed in hepatocytes.^9^ In contrast to semaglutide, HSG4112 can be orally administered. Therefore, HSG4112 is a promising therapeutic for NASH and directly protects hepatocytes.

Lipidomic studies have demonstrated that NASH is associated with a marked alteration in the hepatic lipidome, which is correlated with the disease progression.^26,27^ The changes in the levels of glycerolipids, phospholipids, and fatty acids are closely associated with the progression of NASH. In particular, NASH is correlated with the downregulation of plasma PS, PE, PI, PC, and SM contents^28,29^ and upregulation of LPLs. The accumulation of LPC, an important mediator of hepatic lipotoxicity, disrupts mitochondrial integrity and promotes the release of pro-inflammatory molecules.^30–32^ Patients with NASH are characterized by increased levels of saturated FAs and decreased levels of polyunsaturated FAs.^33,34^ Unsaturated FAs mitigate the saturated FFA-induced hepatic lipotoxicity.^35–37^ Thus, lipidome profile analysis can aid in evaluating the therapeutic efficacy of drugs against NASH. In this study, the levels of individual lipid species in the lipidome were markedly different between the HSG4112-treated and semaglutide-treated groups. HSG4112 upregulated SM and PC species, whereas semaglutide upregulated only some PC species but did not affect the levels of SM species. In this study, the dissipation of LPC, LPE, and LPG in the HSG4112-treated group was markedly higher than that in the semaglutide-treated group. Additionally, HSG4112 downregulated most FFA species, especially glycerolipids. Thus, these effects of HSG4112 may be closely associated with the alleviation of hepatic steatosis and inflammation. While semaglutide promoted the accumulation of some saturated FAs in the liver, it also increased the levels of various unsaturated FAs. Therefore, semaglutide may also effectively alleviate FFA-induced hepatic lipotoxicity. However, further studies are needed to examine the potential side effects of semaglutide-induced upregulation of LPLs. These findings indicate that HSG4112 protects against hepatic lipotoxicity by altering the levels of particular lipid species in the hepatic lipidome.

Autophagy is important for the maintenance of liver homeostasis.^38^ Metabolic disorders are characterized by markedly decreased autophagy activity, which leads to enhanced lipid accumulation, mitochondrial dysfunction, oxidative stress, and inflammation.^39–41^ In particular, autophagy promotes lipid catabolism by breaking down lipid droplets and protecting the hepatocytes from inflammation and oxidative stress.^42–44^ Transcriptome and proteome data demonstrated that HSG4112 activated the autophagic pathways (autophagy initiation, autophagy/mitophagy assembly, and autophagy maturation). HSG4112 markedly downregulated lipid accumulation through autophagy activation in hepatocytes. In NAFLD, the overloading of saturated FFAs into mitochondria promotes mitochondrial dysfunction through the accumulation of ROS in hepatocytes, which exacerbates lipotoxicity and inflammation in the liver. The levels of free radicals generated from FAO are higher than those generated from glucose catabolism under lipotoxic conditions.^15,45^ Consistent with these previous findings, PA treatment increased FAO and enhanced the production of mitochondrial superoxide in hepatocytes in this study. However, HSG4112 decreased oxidative stress to the level observed in the control, upregulated FAO, and decreased lipid peroxidation. These results suggest that HSG4112 alleviates oxidative stress and promotes lipid catabolism by inducing autophagy. Mitophagy clears damaged mitochondria resulting from oxidative stress under pathological conditions.^46–49^ Additionally, mitophagy maintains mitochondrial homeostasis under physiological conditions.^50^ HSG4112 induced mitophagy activation and alleviated oxidative stress, which indicated that it can induce mitophagy independent of oxidative stress. The phospho-proteome data demonstrated that HSG4112 induces autophagy/mitophagy activation through AMPK signaling, which is the major signaling pathway for mitophagy activation in metabolic diseases.^45,51^ Thus, HSG4112 alleviates NASH severity, hepatic steatosis, oxidative stress, and inflammation by activating autophagy.

The identification of the drug target is essential to understand the MOA and determine the clinical dose regimen based on the drug-target engagement. Chemical-protein interactome data demonstrated that Pon2 was a specific target protein of HSG4112. Pon2 is not reported to be associated with NASH in clinical studies. However, Pon2 has a critical role in the pathogenesis of NASH in animal models.^52,53^ In the liver of the NASH rat model, Pon2 expression is upregulated and PON2 activity is downregulated through oxidative inactivation.^52^ In the NASH mouse model, PON2 deficiency decreases energy expenditure and oxidative capacity, which are etiological factors for obesity, an important risk factor for NASH.^53^ Thus, PON2 is a potential therapeutic target for NASH.

The PON family comprises the following three members: PON1, PON2, and PON3.^54^ PON2, which exhibits esterase and lactonase activities, is expressed in most tissues, including the liver.^55,56^ Additionally, PON2 is primarily localized to the mitochondria and endoplasmic reticulum, which are major sites of oxidative stress, and exert antioxidant and anti-inflammatory effects.^57,58^ Therefore, PON2 deficiency promotes mitochondrial dysfunction by increasing oxidative stress.^19^ Mitochondrial localization of HSG4112 indicates that it may be closely associated with the mitochondrial Pon2. HSG4112 promotes Pon2 activity without affecting its expression. Additionally, HSG4112 upregulates FAO and decreases the production of mitochondrial superoxide through Pon2. Furthermore, the antioxidant effect of HSG4112, which downregulated total lipid peroxidation, is dependent on Pon2 activity. Therefore, HSG4112 may exert therapeutic effects by promoting Pon2 activity. *Pon2* KD was hypothesized to induce autophagy/mitophagy activation as it increased oxidative stress in hepatocytes. However, *Pon2* KD inhibited autophagy/mitophagy under basal and lipotoxic conditions. HSG4112-induced autophagy/mitophagy activation was diminished in *Pon2* KD cells, which led to lipid accumulation. Thus, HSG4112 alleviates NASH through Pon2 and by activating autophagy/mitophagy, promoting lipid metabolism, and maintaining redox homeostasis.

In summary, these findings indicate that HSG4112-mediated alleviation of hepatic injury was correlated with increased FAO and autophagy and decreased lipid accumulation, inflammation, and mitochondrial oxidative stress. Mechanistically, the therapeutic effects of HSG4112 were dependent on Pon2 activity. Hepatic Pon2 activation by HSG4112 is closely associated with autophagy activation and antioxidant effects. Thus, HSG4112 is a pharmacological activator of Pon2 and a potential therapeutic for NASH.

## METHODS

### Animals

All animal experiments were performed according to the internationally accepted principles for the care and use of laboratory animals. The license for performing animal experiments was issued by the Danish Committee for animal research for Jacob Jelsing (2013-15-2934-00784). Male C57BL/6J mice aged 5 weeks (Janvier Labs, France) were housed in a controlled environment (circadian cycle, 12-h light/dark cycle (lights on at 3 AM); temperature, 21 ± 2°C; humidity, 50% ± 10%). Individual mice were tagged with an implantable microchip (PetID Microchip, E-vet, Haderslev, Denmark). The animals had ad libitum access to tap water and a diet comprising 40% fat (18% trans-fat), 40% carbohydrates (20% fructose), and 2% cholesterol (D09100301, Research Diets, USA). Mice were fed on this diet for 35–37 weeks to establish a severe NASH phenotype. The liver of all mice was biopsied before drug treatment. Fibrosis and steatosis stages in the liver biopsy were analyzed at baseline (fibrosis score ≥ 1, steatosis score ≥ 2). Individual animals were placed in separate cages. The total number of animals used for each experiment is indicated in the figures and table. An outline of the study design is shown in Figures 1 and 2.

### Baseline liver biopsy

The liver was biopsied 4 weeks before drug treatment. Briefly, mice were pretreated with enrofloxacin (Bayer, Germany) one day before biopsy. Before the biopsy, mice were anesthetized with isoflurane in atmospheric air. A midline abdominal incision was introduced to expose the left lateral lobe and a cone-shaped liver biopsy sample (50–100 mg) was collected. The sample was fixed in 4% paraformaldehyde overnight and subjected to histological analysis. The cut surfaces were electrocoagulated using an electrosurgical unit (ERBE VIO 100C, ERBE, GA, USA). The liver was returned to the abdominal cavity, the abdominal wall was sutured, and the skin was stapled. Mice were intraperitoneally administered with carprofen (Pfizer, USA) (5–0.01 mL/kg bodyweight) at the time of surgery and on days 1 and 2 post-surgery to manage postoperative pain and infection. At week 3 post-biopsy, mice were randomized into various treatment groups based on bodyweight and liver fibrosis and steatosis scores.

### Drug treatment

SGDs were prepared at Glaceum Inc. (Suwon, Republic of Korea) following the protocols mentioned in the patent US9783551B2. Mice were orally administered with SGDs at doses of 50 and 100 mg/kg bodyweight. One group of mice was subcutaneously administered with semaglutide (30 nmol/kg bodyweight) once daily. Drug treatment and diet were continued for 6 weeks (for experiments comparing the effects of HSG4112 and HSG4113) or 10 weeks (for experiments comparing the effects of semaglutide and HSG4112). Food intake was measured once daily for 2 weeks after the initiation of drug treatment. The blood samples were collected from the tail vein of mice under fasting conditions five days before the end of the experiment to determine the blood glucose and insulin levels. Additionally, a terminal blood sample was collected from the tail vein of non-fasting mice for plasma biochemical analysis. Mice were sacrificed by cardiac puncture under isoflurane anesthesia. The liver samples were excised, weighed, and processed for further analysis.

### Bodyweight and body composition analysis

Bodyweight was monitored daily during the treatment period. Whole-body fat and lean mass were analyzed during the 10 weeks of treatment using non-invasive EchoMRI scanning with EchoMRI-900 (EchoMRI, TX, USA). During the scanning procedure, mice were placed in a restrainer for 90–120 s. To analyze the body composition parameters at the end of the study period, the weights of the liver, epididymal white adipose tissue (EWAT), peri-renal white adipose tissue, and gastrocnemius muscle were determined.

### Plasma and tissue biochemical analyses

#### Evaluation of Fgf21, Lep, and cytokines in the cardiac blood samples

The abdominal cavity of mice was opened under isoflurane anesthesia and the cardiac blood sample was collected using a syringe. The blood sample was transferred into a microvette/vacuette containing an anticoagulant, mixed five times by inversion, stored at 4°C until centrifugation. The plasma samples (supernatant) were transferred to new tubes and immediately frozen on dry ice and stored at −80°C until use.

#### Evaluation of glucose, insulin, ALT, AST, ALP, and total cholesterol (TC) levels in the tail blood samples

Tail blood (≤ 200 µL) was pressed into an open microvette (200 µL) containing an appropriate anticoagulant. The blood samples were stored at 4°C until centrifugation. The plasma was separated, transferred to new tubes, and immediately frozen on dry ice and stored at −80°C until use.

#### Evaluation of blood glucose and insulin levels

The blood samples were collected into heparinized glass capillary tubes and immediately suspended in glucose/lactate system solution buffer (EKF-Diagnostics, Germany). Next, the blood glucose levels were measured using a BIOSEN c-Line glucose meter (EKF-Diagnostics), following the manufacturer’s instructions. For analyzing insulin levels, the blood samples were collected in heparinized tubes and the plasma was separated. The plasma samples were stored at −80°C until use. The insulin levels were measured using the MSD platform (Meso Scale Diagnostics), following the manufacturer’s instructions.

#### Evaluation of TC, ALT, AST, and ALP

The blood samples were collected in heparinized tubes and the plasma was separated. The plasma samples were stored at −80°C until use. The TC, ALT, AST, and ALP levels were measured using commercial kits (Roche Diagnostics) with the Cobas c501 autoanalyzer, following the manufacturer’s instructions.

#### Evaluation of Fgf21, cytokines, and Lep

The blood samples were collected in ethylenediaminetetraacetic acid (EDTA)-coated tubes and the plasma was separated. The plasma samples were stored at −80°C until use. The Fgf21, Tnfa, Il10, Il5, Il6, and Cxcl1 levels in the plasma were measured in duplicates using an enzyme-linked immunosorbent assay (ELISA) kit (BioVendor), following the manufacturer’s instructions. The levels of leptin and insulin were measured using the MSD platform (Meso Scale Diagnostics), following the manufacturer’s instructions.

#### Liver TC analysis

The liver samples were homogenized and heated twice at 90°C in 5% NP-40 to extract TC. The samples were centrifuged and the TC content in the supernatant was measured using commercial kits (Roche Diagnostics) with the Cobas c501 autoanalyzer, following the manufacturer’s instructions.

### Liver histological and quantitative analyses

Biopsy samples obtained at the baseline and the end of the study period (both from the left lateral lobe) were fixed overnight in 4% paraformaldehyde. The liver tissue was paraffin-embedded and sectioned (3 µm thickness). The sections were stained with H&E and anti-Lgals3, anti-Acta1, anti-Adgre1, anti-Map1lc3b, and anti-Becn1 antibodies, following the manufacturer’s instructions. The samples were subjected to quantitative histomorphometry using Visiomorph**â** digital imaging software (Visiopharm, Hørsholm, Denmark) or NIH ImageJ software (http://rsbweb.nih.gov/ij/). The primary antibodies used in this study are listed in Supplementary Table 3. The fractional area of liver fat accumulation (macrosteatosis) was determined using HE-stained sections and expressed as a percentage of the total sectional area. The hepatic lipid droplet accumulation, which indicates hepatic steatosis, was confirmed using oil red O staining. Liver fibrosis was examined using picrosirius red staining. The Lgals3-positive and Acta1-positive areas were expressed as a percentage of the total parenchymal area by subtracting the corresponding fat area determined on adjacent HE-stained sections. Macrophage infiltration in the liver was assessed based on Adgre1 immunostaining and Ccl2 expression. Autophagy activation in the liver was determined based on Map1lc3b and Becn1 immunostaining. The staining intensities of Map1lc3b and Becn1 are represented as inverted median pixel values (IMPVs). Hepatic oxidative stress was assessed using the thiobarbituric acid reactive substance (TBARS) assay. To examine adiposity, the paraffin-embedded EWAT sections were subjected to H&E staining. The average adipocyte size in EWAT was measured using morphometry with NIH ImageJ software. All histological assessments were performed by experienced histologists who were blinded to the experimental groups. Steatosis, lobular inflammation, hepatocyte ballooning, and fibrosis in the liver were scored before and after treatment using the NAS and fibrosis staging system.

### RNAseq and data analysis

The liver samples after treatment (15 mg fresh tissue) and gastrocnemius muscle were subjected to RNAseq. The RNA quantity and quality were measured using Qubit^®^ (Thermo Fisher Scientific, OR, USA) and a bioanalyzer with RNA 6000 Nano kit (Agilent, Waldbronn, Germany), respectively. The RNA sequence libraries were prepared using NeoPrep (Illumina, CA, USA) with Illumina TruSeq stranded mRNA Library kit for NeoPrep (Illumina) and sequenced on the NextSeq 500 (Illumina) with NSQ 500 hi-Output KT v2 (75 CTS, Illumina). The sequencing reads were aligned to the GRCm38 v84 Ensembl *Mus musculus* genome using STAR v.2.5.2a with default parameters. Differential gene expression analysis was performed with DEseq2. The difference in the expression of genes with a Benjamini-Hochberg-adjusted p-value of ≤ 0.05 (5% false discovery rate (FDR)) was considered significant.

### Proteomics

#### Protein extraction and digestion

Cryo-pulverized liver tissues (wet weight: 40– 50 mg) were homogenized in 1 mL radioimmunoprecipitation assay (RIPA) lysis buffer (Pierce) (5% sodium dodecyl sulfate (SDS), 0.1 M Tris-HCl (pH 7.6), 1× Halt protease inhibitor (Pierce), and 1× phosphatase inhibitor (PhosSTOP, Roche)). The lysates were centrifuged at 20,000 g for 10 min and the supernatant was transferred to a new tube. The protein concentration in the supernatant was measured using the bicinchoninic acid (BCA) protein assay kit (Pierce). Protein samples (600 μg) were divided into two equal portions, and each portion was digested separately using an S-Trap method (Katelyn et al., 2018). Briefly, 300 μg proteins were incubated with 20 mM dithiothreitol (DTT; Sigma Aldrich) for 10 min at 95°C on a thermomixer (Thermo Fisher Scientific). The proteins were then alkylated with 50 mM iodoacetamide (Sigma Aldrich) for 30 min in the dark. Before digestion, phosphoric acid at a final concentration of 1.2% and six volumes of binding buffer (90% methanol; 100 mM triethylammonium bicarbonate (TEAB); pH 7.1) were added to the sample and gently mixed. The protein solution was loaded to an S-Trap filter and centrifuged at 3000 rpm for 1 min. The flow-through was collected and reloaded onto a filter. This step was repeated thrice. The filter was washed thrice with 300 μL of binding buffer. Digestion was performed with Lys-C (Pierce) for 1 h and trypsin (Pierce) overnight (enzyme-to-protein ratio = 1:50). The peptides were eluted twice using the following three buffers (200 μL each): 50 mM TEAB, 0.2% formic acid in H_2_O, and 50% acetonitrile and 0.2% formic acid in H_2_O.

#### Tandem mass tag (TMT)-16 labeling and peptide fractionation

The eluted peptides (400 µg) from each sample were labeled with 16-plex TMT reagents (Thermo Fisher Scientific, Germany), following the manufacturer’s instructions. The peptides were solubilized in 100 μL of 100 mM TEAB (pH 8.5) solution and incubated with labeling reagent in 20 μL of acetonitrile for 1 h with shaking. To quench the unreacted TMT reagents, 8 μL of 5% hydroxylamine was added. Differentially labeled peptides were then mixed (16 × 400 μg) and dried using a vacuum centrifuge. The quenched, combined sample was desalted using a C18 column (Harvard apparatus). The pooled TMT-labeled peptide sample was fractionated using basic pH reverse-phase liquid chromatography with an Agilent 1260 Infinity HPLC system (Agilent, CA, USA). The chromatography conditions were as follows: column, Xbridge C18 analytical column (4.6 mm × 250 mm); solvent A, 10 mM ammonium formate in water (pH 9.5); solvent B, 10 mM ammonium formate in 90 % acetonitrile (pH 9.5); gradient time, 105 min; flow rate, 500 mL/min. The gradient conditions were follows: 0% solvent B for 10 min; 0%– 13% solvent B for 10 min; 13%–40% solvent B for 60 min; 40%–70% solvent B for 15 min; 70% solvent B for 10 min; 70%–0% solvent B for 10 min. In total, 96 fractions were collected every minute from 15 min to 110 min. The fractions were pooled into 24 non-contiguously concatenated peptide fractions. The resultant 24 fractions were dried and stored at −80°C.

#### Phosphopeptide enrichment

The phosphopeptides from 12 fractions obtained by combining sequential fractions of the 24 fractions (e.g. F1 and F2 or F3 and F4) were enriched using the immobilized metal affinity chromatography (IMAC). IMAC beads were prepared from Ni-nitrilotriacetic acid (NTA) magnetic agarose beads. Ni-NTA beads (500 mL) were washed thrice with water. The beads were then incubated with 100 mM EDTA (pH 8.0) for 30 min with end-over-end rotation to remove nickel ions. The EDTA solution was removed and the beads were washed thrice with water. The NTA beads were treated with 10 mM of aqueous FeCl_3_ solution for 30 min with end-over-end rotation. Iron-chelated IMAC beads were washed thrice with water. Ni-NTA agarose beads were used to prepare Fe^3+^-NTA agarose beads. Approximately 300 μg of peptides from each phosphoproteome fraction were reconstituted in 500 μL of 80% MeCN/0.1% trifluoroacetic acid (TFA). The solution was incubated with IMAC beads for 30 min on a shaker at room temperature. The samples were briefly centrifuged using a tabletop centrifuge. The clarified peptide flow-throughs were separated from the beads and the beads were reconstituted in 200 mL IMAC binding/wash buffer (80 MeCN/0.1% TFA) and loaded onto equilibrated Empore C18 silica-packed stage tips (3M, 2315). The samples were then washed twice with 50 μL of IMAC binding/wash buffer and once with 50 μL of 1% FA and eluted thrice from the IMAC beads to the stage tips with 70 μL of 500 mM dibasic sodium phosphate (pH 7.0, Sigma Aldrich, S9763). The stage tips were then washed once with 100 mL 1% FA. Phosphopeptides were eluted from the stage tips with 60 μL of 50% MeCN/0.1% FA.

#### LC-ESI-MS/MS analysis of the proteome

All peptide samples were separated using an ultra-performance liquid chromatography system equipped with analytical columns (75 µm × 50 cm, C18, 3 µm, 100 Å) and trap columns (75 µm × 2 cm, C18, 3 µm, 100 Å). The mobile phases were as follows: solvent A, 0.1% formic acid in water; solvent B, 0.1% formic acid in 90% acetonitrile. The solvent gradient conditions for global proteome profiling analysis were as follows: 2% solvent B for 8 min, 2%–10% solvent B for 3 min, 10%–25% solvent B for 113 min, 25%–40% solvent B for 20 min, 70% solvent B for 4 min, and 2% solvent B for 20 min. Meanwhile, the gradient conditions for phosphoproteome analysis were as follows: 2% solvent B for 8 min, 2%– 10% solvent B for 3 min, 10%–20% solvent B for 123 min, 20%–30% solvent B for 20 min, 70% solvent B for 4 min, and 2% solvent B for 20 min. For global peptide analysis, 2 µg of peptides from each of the 24 fractions were individually analyzed. All enriched peptides from each of the 12 fractions were injected for phosphopeptide analysis. The eluted peptides from LC were analyzed with a Q exactive plus orbitrap mass spectrometer (Thermo Fisher Scientific) equipped with an easy-spray nano source. Data acquisition was performed using Xcalibur Q Exactive v2.1 software in positive ion mode at a spray voltage of 2.0 kV. MS1 spectra were captured under the following conditions: resolution, 70,000; AGC target, 1e6; mass range, 400–1800 m/z; mode cycle time, 2 s. MS2 spectra were captured under the following conditions: resolution, 35,000; AGC target, 2e5; isolation window, 2.0 m/z; maximum injection time, 100 ms; HCD collision energy, 32%. The peptide mode was selected for monoisotopic peak determination. Charge state screening was enabled to only include precursor charge states 2–6 with an intensity threshold of 5e4. Peptides that triggered MS/MS scans were dynamically excluded from further MS/MS scans for 30 s with a mass tolerance of 10 ppm (±).

#### Protein and phosphopeptide identification

The acquired raw files were processed using Proteome Discoverer v.2.4 (Thermo Fisher Scientific) software environment with the built-in Sequest HT search engine for the identification of peptides and phosphopeptides. The data were searched using a target-decoy approach against Uniprot Human (Jan 2021, 20,452 entries) reference proteome (FASTA file) (FDR < 1%) at the level of proteins, peptides, and modifications using minor changes to the default settings as follows: oxidized methionine (M); in case of phosphopeptides, search phospho (S,T,Y) was selected as variable modifications, while carbamidomethyl (C) selected as fixed modification. A maximum of 2 missed cleavages and a minimum peptide length of seven amino acids were allowed. Enzyme specificity was set to trypsin. An initial precursor mass deviation up to 10 ppm and a fragment mass deviation up to 0.06 Da were allowed. To quantify each reporter ion in the sample, ‘reporter ion quantifier’ with TMT 16-plex was used. For highly confident quantifications of protein, the protein ratios were calculated from two or more unique quantitative peptides in each replicate. All reporter ion intensities were transformed to log2 values. Proteins that did not display all values in at least one group were filtered out.

#### Proteomic data analysis

For global proteome analysis, the TMT intensities of the peptides were normalized using the quantile normalization method. Differentially expressed proteins (DEPs) were identified using an integrative statistical method. Briefly, log2 (intensity) of each protein was applied to its abundance for all replicates. The fold change and adjusted p-value were calculated using Student’s t-test. Finally, DEPs that had a combined fold change value of 1.5 and a p-value < 0.05 were selected. For phosphoproteome analysis, the same normalization and statistical method described above were used. Differentially phosphorylated peptides (DPPs) that were uniquely assigned to the protein were selected. To explore the signal pathway in which the DEPs and DPPs are enriched, Kyoto Encyclopedia of Genes and Genomes (KEGG) enrichment and Reactome pathway analyses were performed using the g:Profiler. To reconstruct drug-related network modeling, protein-protein interactions (PPIs) were obtained from the STRING database. Network models were developed for a list of the selected proteins for network analysis based on the collected PPIs. In the network models, the nodes were arranged based on the KEGG and Reactome Pathway database analyses.

### Lipidomics

#### Lipid extraction

Liver samples were weighed and individually cryo-pulverized using a Cryoprep device (CP02, Covaris). The lyophilized samples from each tissue were aliquoted to separate tubes (50 mg for proteomic analysis and 20 mg for lipidomic analysis) and stored at −80°C. For lipid analysis of liver tissues, two-step lipid extraction method was performed. To perform the neutral extraction, liver tissues were thawed on ice and incubated with internal standards (10 μL) and methanol/chloroform (2:1 (v/v); 990 μL). The samples were vortexed for 60 s. The standard mix SPLASH Lipidomix Mass Spec Standard j 330707 supplemented with LPG (14:0), LPI (13:0), LPA (14:0), Cer (d18:1−12:0), So (d17:1), Sa (d17:0), CerP (d18:1−12:0), So1P (d17:1), Sa1P (d17:0), and FFA (20:4-d8) was used. The sample was incubated for 10 min on ice and centrifuged 15,000 *g* and 4°C for 2 min. Next, 950 μL of supernatant was transferred to a new tube. To perform the acidic extraction, the remaining tissue was incubated with 750 μL of chloroform/methanol/37% HCl (40:80:1 v/v/v) mixture for 15 min at room temperature. Next, 250 μL of cold chloroform and 450 μL of cold 0.1 N HCl were added to the sample. The samples were vortexed for 1 min and centrifuged at 7000 g and 4°C for 2 min. The bottom organic phase was collected and pooled with a prior extract and dried using a SpeedVac concentrator.

#### Lipid measurement using LC-MS

HPLC-ESI-MS/MS analyses were performed using a 1200 series HPLC system (Agilent Technologies, DE, USA) coupled to 6490 Accurate-Mass Triple Quadrupole Mass Spectrometer (Agilent Technologies). The LC-MS/MS conditions were as follows: column, Hypersil GOLD column (2.1 × 100 mm ID; 1.9 μm, Thermo Fisher Scientific); solvent A (acetonitrile/methanol/water (19:19:2 v/v/v) + 20 mmol/L ammonium formate + 0.1% (v/v) formic acid); solvent B, (2-propanol +20 mmol/L ammonium formate + 0.1% (v/v) formic acid); flow rate, 250 μL/ min; total run time, 33 min; sample injection volume, 5 μL. The gradient elution program was as follows: 0–5 min, solvent B 3%; 5–18 min, solvent B 5–30%; 18–24 min, solvent B 30%–90%; 24–28 min, solvent B 90%; 28–29 min, 90%–3%; and 29–33 min, solvent B 3%. The parameters of operating source conditions were as follows: capillary voltage in positive mode, 3500 V; capillary voltage in negative mode, 3000 V; sheath gas flow rate, 11 L/min (UHP nitrogen); sheath gas temperature, 200°C; drying gas flow rate, 15 L/min; drying gas temperature, 150°C; nebulizer gas pressure, 25 psi. The optimal dMRM conditions were used to analyze various lipid species.

#### Lipidomics data analysis

Agilent Mass Hunter Workstation Data Acquisition software was used to process the LC/MS data. Qualitative Analysis B.06.00 software (Agilent Technologies) was used to export the m/z of precursor and product ions and retention time of target lipids in the multiple reaction monitoring data. The area of the assigned peak in the raw data was calculated using an in-house database constructed using Skyline software package (MacCoss Laboratory, University of Washington, WA, USA). Lipid abundance was normalized to tissue weight and internal standard peak area. MetaboAnalyst Web site (https://metaboanalyst.ca) was used to perform PCA and projection to latent structure discriminant analysis (PLS-DA). Unweighted correlation network graphs were generated based on Spearman’s correlation. All nodes with P values less than 0.05 were used in SPSS version 21.0 (IBM Corp, NY, USA). Heatmaps were generated using the Morpheus Web site (https://software.broadinstitute.org/morpheus).

### Cell culture and generation of stable *Pon2* knockdown cells

L02 cells (an immortalized normal liver cell line) were a kind gift from Dr. KH Lee (Korea Institute of Radiological and Medical Sciences). The cells were cultured in Dulbecco’s modified Eagle’s medium (DMEM) (Welgene, LM001-05) supplemented with 10% fetal bovine serum and 100 U/mL of penicillin and streptomycin at 37°C in a 5% CO_2_ incubator.

Stable *Pon2* knockdown (KD) L02 cells were generated by transducing the cells with recombinant lentivirus harboring short hairpin RNA against *Pon2* (shPON2; Gencopoeia, LPP-HSH013480-LVRU6P). Pon2 KD cells were cultured in DMEM supplemented with puromycin (500 ng/mL) (Life Technologies, A11138-03). The KD of *PON2* was confirmed using immunoblotting with the anti-Pon2 antibodies.

Cells stably expressing mt-Keima were generated by transducing the cells with recombinant lentivirus containing mt-Keima cDNA for 48 h. The recombinant cells were selected with puromycin (500 ng/mL). The packaging vectors pSPAX2 and pMD2.0G were co-transfected with pmt-Keima into HEK293T cells for 72 h. The cell supernatant was filtered through a 0.22-µm membrane filter. The virus was concentrated by subjecting the samples to ultra-centrifugation at 20000 rpm. The supernatant was discarded and the viruses were suspended in an appropriate amount of PBS and stored at −80°C until use.

Bovine serum albubim (BSA)-conjugated oleic acid (OA, O3008) and PA (P0500) were purchased from Sigma Aldrich (St. Louis, MO). To prepare the BSA-conjugated PA solution, 100 mM PA solution was prepared in 0.1 mM NaOH and the solution was heated at 70°C. PA solution was incubated with 10% BSA at 55°C for 30 min to obtain 5 mM PA/1% BSA. The solution was then cooled to 25°C, filter-sterilized, and stored at −20°C until use. The cells were incubated with culture medium containing 400 µM FFA (150 µM BSA-conjugated OA and 250 µM BSA-conjugated PA) in the presence or absence of 3 µM HSG4112.

### Lipid accumulation assay

Cells were plated in six-well plates and incubated with 400 µM FFA in the presence or absence of 3 µM HSG4112 for 72 h. The cells fixed with 4% paraformaldehyde for 10 min were stained with oil red O solution (Sigma Aldrich, O1391) for 15 min at room temperature. After washing once with 60% isopropanol, the cells were rinsed with distilled water. The images were captured using an inverted microscope (Carl Zeiss Axioimager M2 fluorescence microscopy). The signal intensity was analyzed using the NIH ImageJ software.

### Autophagy flux analysis

Immunoblotting analysis was performed to analyze the endogenous LC3B-II/I and Sqstm1/p62 expression levels with anti-Map1lc3b and anti-Sqstm1 antibodies, respectively, following the manufacturer’s instructions (Supplemental Table 3). The cells were plated in six-well plates and incubated with 400 µM FFA in the presence or absence of 3 µM HSG4112 for the indicated durations. Next, the cells were lysed with SDS lysis buffer supplemented with a protease inhibitor cocktail (Thermo Fisher Scientific, 78441). The proteins were subjected to immunoblotting. The intensity of each protein signal was normalized to that of Actb. The control value was set to 1.0 and the protein intensity was represented relative to that of the control.

To monitor the autophagy level, LC3B puncta in cells transfected with mRFP-GFP tandem fluorescent-tagged LC3B (tfLC3B) were examined using immunofluorescence microscopy. The cells cultured on glass coverslips were transfected with mRFP-GFP tfLC3B plasmid. At 16 h post-transfection, the cells were treated with 400 µM FFA and 3 µM HSG4112 in the presence or absence of bafilomycin A1 (Invivogen, tlrl-baf1). Cellular localization of LC3B was observed using a Carl Zeiss Confocal LSM710 Meta microscope and the images were processed with the software supplied by the manufacturer (Carl Zeiss, Seoul, Republic of Korea) and analyzed with NIH ImageJ software. Cells containing three or more mRFP-LC3B puncta were defined as autophagy-positive cells. The percentage of autophagy-positive cells relative to the total number of mRFP-positive cells was calculated. At least 100 mRFP-positive cells per sample were counted in at least three independent experiments. Cells stained with both RFP and GFP were defined as autophagosome-positive, while those stained with RFP only were defined as autolysosome-positive. The number of fluorescent LC3B puncta was determined by counting more than 100 cells with triplicates.

### Mitophagy measurement

Endogenous Prkn, Pink1, Bnip, Bnip3l, Map1lc3b, and Sqstm1 expression levels were analyzed using immunoblotting analysis with the respective antibodies, following the manufacturer’s instructions (Supplemental Table 3). The cells were plated into six-well plates and incubated with 125 µM PA in the presence or absence of 3 µM HSG4112 for 24 h. Next, the cells were lysed with SDS lysis buffer supplemented with a protease inhibitor cocktail (Thermo Fisher Scientific, 78441). Immunoblotting was performed as described above.

To monitor the mitophagy level, an mt-Keima-based mitophagy assay was performed using immunofluorescence microscopy. The cells expressing mt-Keima were plated onto a confocal dish, incubated with 125 µM PA in the presence or absence of 3 µM HSG4112 for 24 h, and analyzed using confocal microscopy. Mt-Keima protein was excited at 458 m (neutral, pseudo-colored in green) and 561 nm (acidic, pseudo-colored in red) and detected through the same emission filter (570–695 nm). Laser power was set up at the lowest output to enable the clear visualization of the mt-Keima signal and individualized for each experimental condition. Imaging settings were maintained with the same parameters for comparison between different experimental conditions. Additionally, the images were acquired using a Carl Zeiss Confocal LSM710 Meta microscope (Carl Zeiss, Seoul, Republic of Korea). The fluorescence intensity was quantified using the NIH ImageJ software.

### Measurement of mitochondrial superoxide

Cells were plated in six-well plates and incubated with 125 µM PA in the presence or absence of 3 µM HSG4112 for 24 h. Next, the cells were pulsed with 2.5 µM MitoSOX red mitochondrial superoxide indicator (Thermo Fisher Scientific, M36008) and 5 µM Hoechst 33342 (Thermo Fisher Scientific, H1399) for 30 min and subjected to live cell imaging at 37°C. After washing, the cells were observed under a confocal microscope in a chamber heated to 37°C at 5% CO_2_. Images were acquired using a Carl Zeiss Confocal LSM710 Meta microscope (Carl Zeiss, Seoul, Republic of Korea). The fluorescence intensity was quantified using the NIH ImageJ software.

### Lipid peroxidation assay

Lipid peroxidation in the liver tissue and cells was determined by measuring the contents of malondialdehyde (MDA) resulting from the thiobarbituric acid reaction. The cells were plated in six-well plates and incubated with 125 µM PA in the presence or absence of 3 µM HSG4112 for 24 h. Next, the cells were trypsinized and centrifuged. The cell pellets were washed with PBS and stored at −80°C until use. The MDA concentration was determined using the TBARS parameter assay kit (R&D systems, KGE013), following the manufacturer’s instructions.

For performing the 4-hydroxynonenal (4-HNE) assay, the cells were lysed using RIPA buffer (Cell Signaling Technology) supplemented with a protease inhibitor cocktail (Thermo Fisher Scientific, 78441). The protein concentration was determined using the BCA protein assay kit (Thermo Fisher Scientific, 23225). The 4-HNE concentration was determined using the 4-HNE ELISA kit (BioVision, E4645-100), following the manufacturer’s instructions.

### Measurement of oxygen consumption rate (OCR)

OCR in human liver cells was measured using XFp Extracellular Flux Analyzers (Agilent Seahorse Biosciences). The cells were plated into XFp cell culture mini plates for 24 h. The culture medium was replaced with XF-Base medium (non-buffered Roswell Park Memorial Institute-1640 medium containing 2 mM L-glutamine, 1 mM sodium pyruvate, and 10 mM glucose, pH 7.4) for 30 min. Next, the cells were incubated with HSG4112 (3 µM) for 15 min, followed by incubation with 250 µM palmitate-BSA or BSA (control). Analysis was performed using the XF assay with the cell mito stress test kit (Agilent Seahorse Biosciences). Three measurements were assessed under basal conditions and after the addition of 2 µM oligomycin, 0.5 µM carbonyl cyanide-p-trifluoromethoxyphenylhydrazon (FCCP), and 1 µM rotenone/antimycin. The cellular protein levels were evaluated using the BCA protein assay kit (Thermo Fisher Scientific, 23225). OCR values were normalized to the protein concentration.

To determine FAO, cells were cultured in XFp cell culture mini plates for 24 h. The culture medium was replaced with substrate limited medium for 30 min. Next, the cells were treated with etomoxir (ETO; 40 µM final concentration, Agilent Seahorse Biosciences) in the presence or absence of HSG4112 (3 µM) for 15 min, followed by incubation with 250 µM palmitate-BSA or BSA (control). Analysis was performed using the XF assay with the cell mito stress test kit (Agilent Seahorse Biosciences). Three measurements were assessed under basal conditions and after the addition of 2 µM oligomycin, 0.5 µM FCCP, and 1 µM rotenone/antimycin. The cellular protein levels were evaluated using the BCA protein assay kit (Thermo Fisher Scientific, 23225). The OCR values were normalized to the protein concentration.

### Intracellular localization of HSG4112

Cells were cultured on glass coverslips and pulsed in the presence of propargyl-HSG4112 (3 µM) for 5 h. After labeling, the cells were washed thrice with PBS and incubated in serum-free media with 100 nM MitoTracker Red (Invitrogen, M7512) for 30 min at 37°C in a 5% CO_2_ incubator. The cells were then fixed with 4% paraformaldehyde in PBS for 10 min and stained with 10 µM Alexa488-azide (Invitrogen, A10266) for 30 min using the Click-iT cell reaction buffer kit (Invitrogen, C10269), following the manufacturer’s instructions. Next, the cells were washed thrice with PBS. The images were acquired using a Carl Zeiss Confocal LSM710 Meta microscope (Carl Zeiss, Seoul, Republic of Korea). The data were analyzed using NIH ImageJ software.

### Quantification of Pon2 enzyme activity

#### Intracellular Pon2 activity

Cells were plated into six-well plates and incubated with 125 µM PA in the presence or absence of 3 µM HSG4112 for 24 h. Next, the cells were trypsinized and centrifuged. The cell pellets were washed with PBS and stored at −80°C until use. The frozen cell pellets were incubated with 25 mM Tris buffer (pH 7.4) containing 0.05% n-dodecyl-β-D-maltoside (Merck, D4641) and 1 mM CaCl_2_, and lysed by subjecting them to three freeze-thaw cycles.

For evaluating the Pon2 esterase activity, p-nitrophenyl acetate (pNPA) hydrolysis was determined using a SpectraMax Plus384 microplate reader (Agilent Technologies, Seoul, Republic of Korea). The cell lysates were transferred to a 96-well plate and reactions (0.2 mL final mixture volume) were initiated by adding 1 mM pNPA in Pon2 activity assay buffer (50 mM Tris (pH 8.0) with 1 mM CaCl_2_). The increase in absorbance at 412 nm resulting from the release of p-nitrophenol was monitored.

For evaluating the Pon2 lactonase activity, the enzymatic hydrolysis of the thioalkyl-substituted lactones was determined. The cell lysates were transferred to a 96-well plate and reactions were initiated by adding 1 mM 5-thiobutyl butyrolactone (TBBL) and 1 mM 5′,5-dithiobis (2-nitrobenzoic acid) (DTNB) (Sigma Aldrich, D8130) in Pon2 activity assay buffer. The enzymatic hydrolysis was monitored by examining the absorbance of the reaction mixture at 420 nm.

#### Recombinant PON2 activity

For performing the oxidized linoleic acid (Ox-LA)-mediated Pon2 inhibition assay, purified recombinant Pon2 protein (10 µM) was incubated with or without HSG4112 (10 µM) in the Pon2 activity assay buffer in a 96-well plate for 10 min at room temperature. To inhibit Pon2 activity, the samples were incubated with Ox-LA (100 µM) for 10 min at RT. For evaluating the esterase activity, the reactions were initiated by adding 1 mM pNPA in Pon2 activity assay buffer. The enzymatic hydrolysis was monitored by measuring the absorbance of the reaction mixture at 412 nm. For evaluating the lactonase activity, the reactions were initiated by adding 1 mM TBBL and 1 mM DTNB in Pon2 activity assay buffer. The enzymatic hydrolysis was monitored by measuring the absorbance of the reaction mixture at 420 nm.

### Chemical-protein interactome analysis for identifying HSG4112 target proteins

#### Preparation of protein samples

Biotin-conjugated HSG4112 was synthesized by incubating 20 µM propargyl-HSG4112 with 10 µM biotin-azide (Invitrogen, B10184) using the Click-iT cell reaction buffer kit (Invitrogen, C10269) for 30 min, following the manufacturer’s instructions. Biotin-conjugated HSG4112 was incubated with Dynabbeads^TM^ M-270 streptavidin (Invitrogen, 65305) at 4°C for 1 h. The HSG4112-bead complex was washed thrice with PBS. The cell lysates were incubated with the HSG4112-bead complex at 4°C overnight. The beads were washed thrice with PBS and the bound proteins were eluted with SDS-PAGE sample buffer and stored at −80°C until use.

#### Interactome analysis of target proteins of HSG4112

To identify the HSG4112-binding proteins, the bound proteins were digested using the previously reported FASP protocol (Jung, et al, 2018). Briefly, the proteins were reduced in SDS lysis buffer (4% (w/v) SDS; 0.1 M Tris/HCl (pH 7.6); 0.1 M DTT) at 37°C for 45 min and boiled at 95°C for 10 min. The solution was concentrated using a 30 k membrane filter (Microcon devices, YM-30, Millipore, MA). The buffer was replaced with 0.2 mL UA solution (8 M urea in 0.1 M Tris/HCl (pH 8.5)). The concentrates were mixed with 0.1 mL of 50 mM indole-3-acetic acid (IAA) in UA solution and incubated in the dark at room temperature for 30 min. The samples were centrifuged, washed with 0.2 mL of 50 mM TEAB, and centrifuged thrice at 14,000 *g* for 30 min. Next, 500 ng of trypsin prepared in 0.1 mL of 100 mM TEAB (with the enzyme-to-protein ratio of 1:50) was added to the filter and the samples were incubated at 37°C overnight. The peptides were collected by centrifuging the filter units at 14,000 *g* for 30 min. TEAB (50 µL, 50 mM) was added to the filter and the samples were centrifuged twice at 14,000 g for 20 min. Analytical columns (75 µm × 50 cm, C18, 3 µm, 100 Å) and trap columns (75 µm × 2 cm, C18, 3 µm, 100 Å) were used to separate the peptide samples. The solvents A and B were 0.1% formic acid in water and 0.1% formic acid in acetonitrile, respectively. The proteome profile analysis conditions were as follows: 2%–30% solvent B for 60 min, 25%–90% solvent B for 2 min, 90% solvent B for 8 min, and 2% solvent B for 20 min. The eluted peptides from LC were subjected to mass spectrometry using a Q Exactive HF-X Mass Spectrometer (Thermo Fisher Scientific, Bremen, Germany). Protein identification was performed using Proteome Discoverer 2.4 (Thermo Fisher Scientific) software and Uniprot Human (Jan 2021, 20,452 entries) database. Cellular components and biological processes in which each protein was enriched were analyzed using GO with g;Profiler.

### Evaluation of HSG4112-Pon2 interaction using the competitive binding assay

To confirm the interaction between HSG4112 and Pon2, the HSG4112-bead complex was prepared as described above and incubated with cell lysates at 4°C overnight. The beads were washed thrice with PBS, eluted with SDS-PAGE sample buffer, and subjected to immunoblotting with the anti-Pon2 antibodies.

For performing the competitive binding assay, cell lysates were incubated with free HSG4112 at the indicated concentrations (0–5 µM) for 5 h at 4°C. Next, the HSG4112-bead complex was incubated with the free HSG4112-treated cell lysates at 4°C overnight. The beads were washed thrice with PBS and the bounded proteins were eluted with SDS-PAGE sample buffer. The samples were then subjected to immunoblotting with the anti-Pon2 antibodies.

### Immunohistochemical analysis of liver tissues

The sections were deparaffinized with xylene and dehydrated with ethanol. Next, the sections were subjected to antigen retrieval and incubated with blocking solution to prevent nonspecific antibody binding. The sections were then probed with primary antibodies at 4°C overnight, following the manufacturer’s instructions (Supplemental Table 3). After counterstaining with hematoxylin QS (Vector Laboratories, H-3404), the sections were dehydrated and mounted. The staining intensity of each protein was measured using the NIH ImageJ software with the IHC Profiler plugin (http://rsb.info.nih.gov/ij/). The Map1lc3b and Becn1 intensities are represented as IMPVs.

### Immunoblotting analysis

To extract proteins, the cultured cells were lysed using SDS lysis buffer (100 mM Tris-HCl, pH 6.8, 10% glycerol, and 1% SDS) supplemented with a protease inhibitor cocktail (Thermo Fisher Scientific, 78441). The protein concentration was determined using the BCA protein assay kit (Thermo Fisher Scientific, 23225). The samples were boiled in 1× sample buffer (10 mM Tris-HCl (pH 6.8), 1% SDS, 5% glycerol, 0.05% bromophenol blue, and 1% β-mercaptoethanol) for 5 min and subjected to SDS-polyacrylamide gel electrophoresis (SDS-PAGE). The resolved proteins were electrotransferred to an Immobilon-P membrane (Merck, IPVH00010). The membrane was probed with specific antibodies listed in Supplementary Table 3. Immunoreactive signals were detected using a LAS-4000 Luminescent Image Analyzer (GE Healthcare Bio-Sciences). The signal intensity was assessed by measuring the relative density of each band and normalizing it to that of Actb using the Multi Gauge software (Fujifilm).

### Statistical analysis

All data, except RNA sequencing data, were analyzed using GraphPad Prism v5.02 software (GraphPad, La Jolla, CA). The data are represented as mean ± standard error unless specified otherwise. The means were compared using the Student’s t-test or two-way analysis of variance (ANOVA), followed by h Bonferroni’s post-hoc test as indicated in each figure legend. The changes in the histological scores before and after treatment were analyzed using the Fisher’s exact test and were compared with those of the vehicle group. All other parameters were analyzed using one-way ANOVA, followed by Dunnett’s post-hoc test. Differences were considered significant at P < 0.05.

## Acknowledgments

This study was supported by the National Research Foundation of Korea (NRF) grants funded by the Korean government (NRF-2020R1A2C3010511, NRF-2021M3A9I2080488, and NRF-2021M3A9H3017086).

## Author Contributions

G.C.S., K.P.K., and K.H.K. designed the experiments and analyzed the data; G.C.S., H.M.L., S.K.Y., D.R., Y.S.P., H.S.P., and N.Y.K. performed the experiments and analyzed the bioinformatics data. All authors discussed the results and contributed to the data analysis; G.C.S. and K.H.K. prepared the manuscript with contributions from all authors.

## Data and materials availability

All data supporting the findings of this study are available within the article and its Supplementary Information files and from the corresponding author upon reasonable request. A reporting summary for this article is available as a Supplementary Information file.

