## supplementary information for "Synthetic glabridin derivatives mitigate steatohepatitis in a diet-induced biopsy-confirmed non-alcoholic steatosis hepatitis mouse model through paraoxonase-2"

**Supplementary Table 1** Summary of the study groups used to evaluate the therapeutic effect of synthetic glabridin derivatives

| Group no. | Group name | Number of animals | Animal model | Dose (mg/kg bodyweight) | Dosing frequency | Dosing method | Study period |
|---|---|---|---|---|---|---|---|
| 1 | Vehicle | 12 | AMLN-NASH | NA | QD | PO | 6 weeks |
| 2 | HSG4112 | 11 | AMLN-NASH | 100 | QD | PO | 6 weeks |
| 3 | HSG4113 | 10 | AMLN-NASH | 200 | QD | PO | 6 weeks |

QD, once daily; PO, per oral; NASH, non-alcoholic steatosis hepatitis; AMLN, amylin; NA, not applicable

**Supplementary Table 2** Summary of the study group used to comparatively evaluate the therapeutic effects of HSG4112 and semaglutide

| Group no. | Group name | Number of animals | Animal model | Dose (mg/kg bodyweight) | Dosing frequency | Dosing method | Study period |
|---|---|---|---|---|---|---|---|
| 1 | Vehicle | 15 | AMLN-NASH | NA | QD | PO | 10 weeks |
| 2 | HSG4112 50 mg/kg bodyweight | 15 | AMLN-NASH | 50 | QD | PO | 10 weeks |
| 3 | HSG4112 100 mg/kg bodyweight | 14 | AMLN-NASH | 100 | QD | PO | 10 weeks |
| 4 | Semaglutide | 14 | AMLN-NASH | 0.12 (30 nM/kg bodyweight) | QD | SC | 10 weeks |

QD, once daily; PO, per oral; SC, subcutaneous; NASH, non-alcoholic steatosis hepatitis; AMLN, amylin; NA, not applicable

**Supplementary Table 3** List of antibodies used in this study

| Name | Source | Identifier |
|---|---|---|
| Anti-Lgals3 | BioLegend | #125402 |

| | | |
|---|---|---|
| Anti-Acta1 | Abcam | ab124964 |
| Anti-Adgre1 | Abcam | ab111101 |
| Anti-Map1lc3b | Cell Signaling Technology | #2775 |
| Anti-Sqstm1/p62 | Cell Signaling Technology | #5114 |
| Anti-Actb | Sigma | A5441 |
| Anti-Map1lc3b (immunohistochemical) | Cell Signaling Technology | #43566 |
| Anti-Becn1 | Atlas antibodies | HPA028949 |
| Anti-Prkn | Cell Signaling Technology | #2132 |
| Anti-Pink1 | Novus | BC100-494 |
| Anti-Bnip3 | Abcam | ab10433 |
| Anti-Bnip3l | Cell Signaling Technology | #12396 |
| Anti-Pon2 | Abcam | ab183718 |
| Anti-mouse IgG, horseradish peroxidase (HRP)-conjugated antibody | Cell Signaling Technology | #7076 |
| Anti-rabbit IgG, HRP linked antibody | Cell Signaling Technology | #7074 |

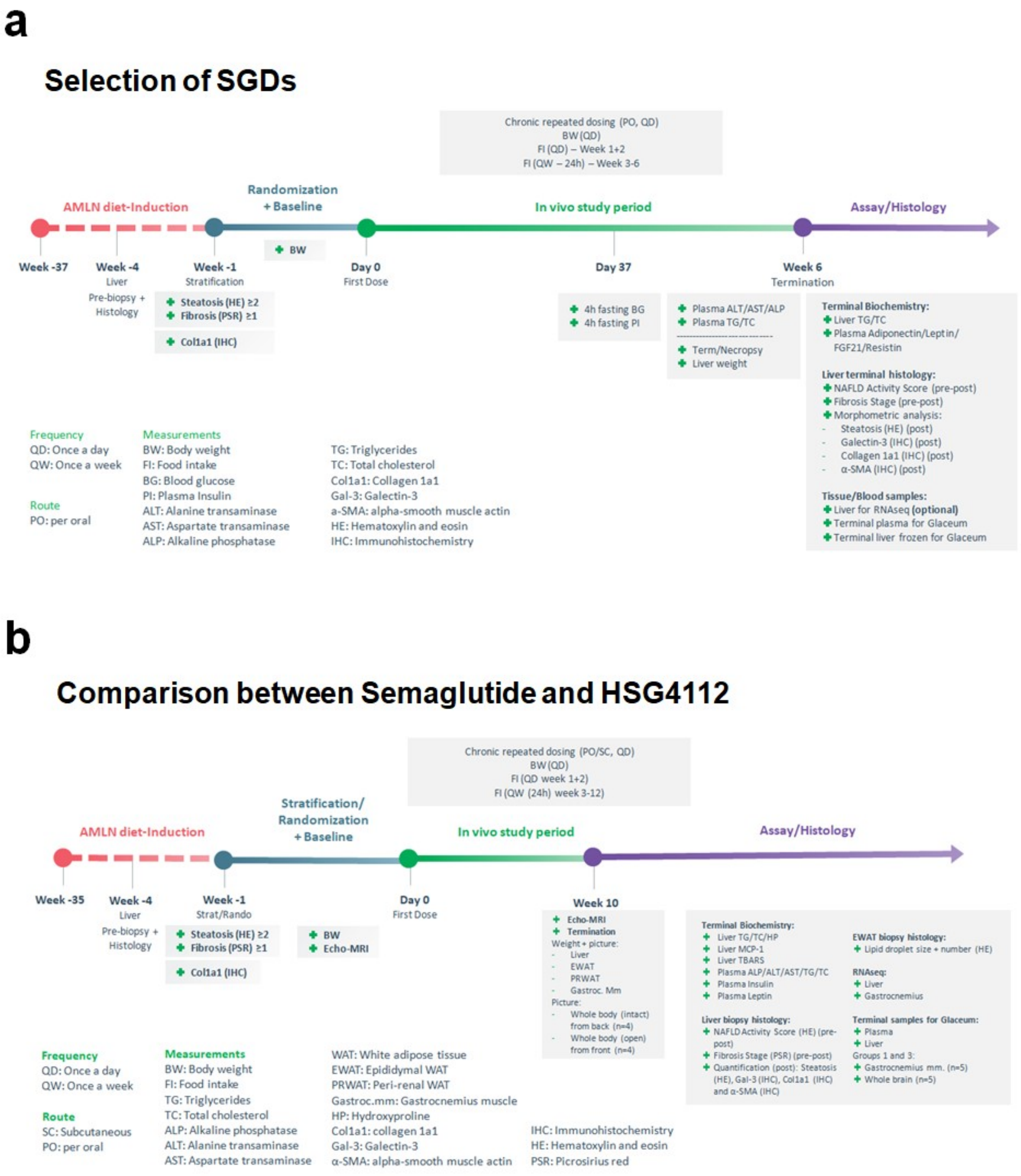


**Supplementary Fig. 1** Summary of the experiments performed to evaluate the therapeutic effects of synthetic glabridin derivatives (SGDs) using the biopsy-confirmed non-alcoholic steatosis hepatitis (NASH) mouse model. **a** Mice were fed an amylin (AMLN) diet for 37 weeks and the mouse liver was biopsied to confirm NASH. Next, the mice were orally administered HSG4112 (100 mg/kg bodyweight) and HSG4113 (200 mg/kg bodyweight) once daily for 6 weeks. **b** Mice were fed with AMLN for 35 weeks and the liver was biopsied to confirm NASH. Next, the mice were administered HSG4112 (50 or 100 mg/kg bodyweight; oral administration) or semaglutide (30 nM/kg bodyweight, subcutaneous injection) once daily for 10 weeks.

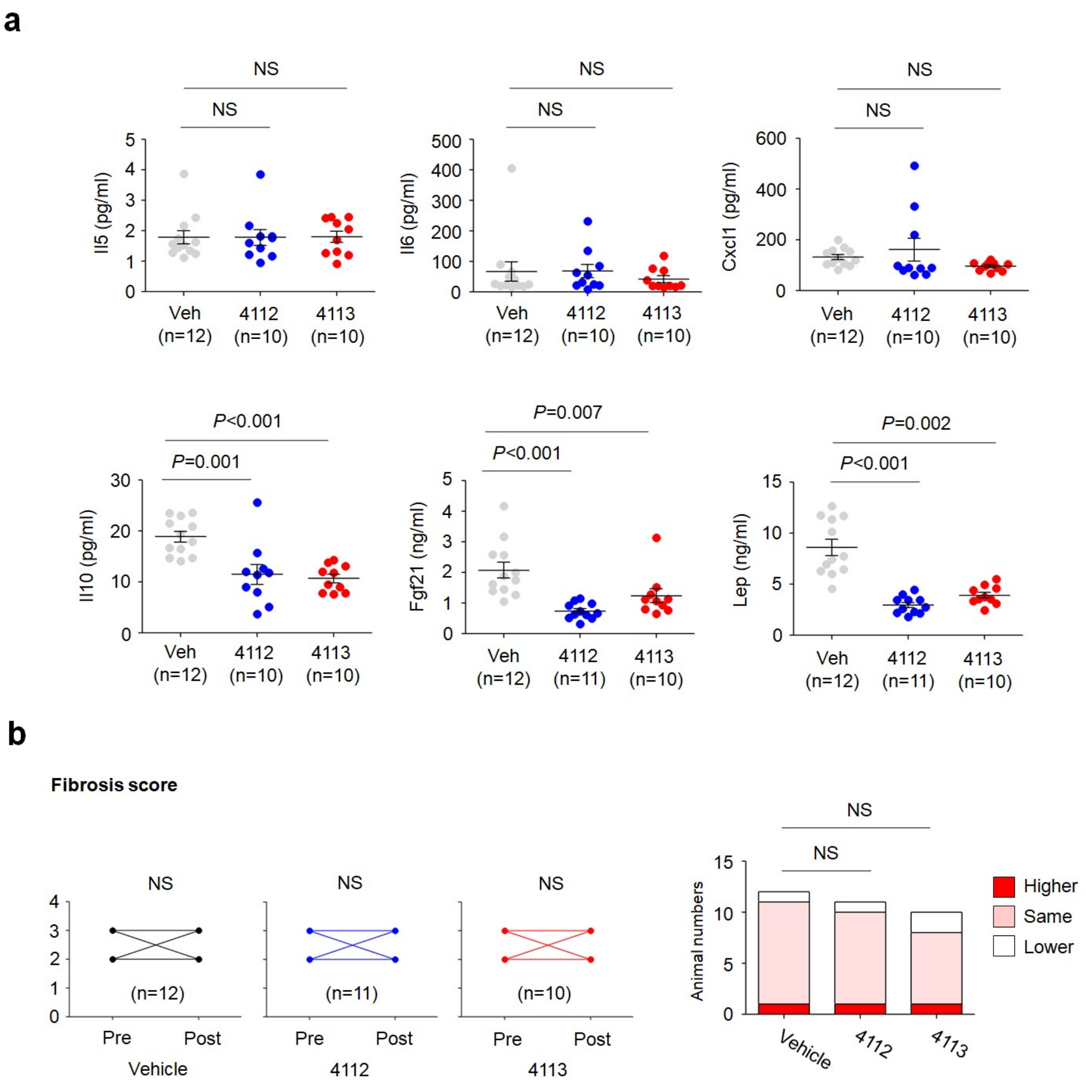


**Supplementary Fig. 2** Histopathological and plasma biochemical parameters in the biopsy-confirmed non-alcoholic steatosis hepatitis model. **a** Quantification of serum levels of Il5, Il6, Cxcl1, Il10, Fgf21, and Lep. **b** Histological scores of fibrosis in biopsies before drug treatment (pre) and at the end of drug treatment (post) are shown on the left. The number of animals exhibiting exacerbation (higher), no change (same), and alleviation (lower) of disease progression after drug treatment relative to pre-treatment is shown on the right.

| Feature | Degree | Score |
|---|---|---|
| Steatosis | <5% | 0 |
| | 5-33% | 1 |
| | >33-66% | 2 |
| | >66% | 3 |
| Lobular inflammation | No foci | 0 |
| | <2 foci/200x | 1 |
| | 2-4 foci/200x | 2 |
| | >4 foci/200x | 3 |
| Ballooning degeneration | None | 0 |
| | Few | 1 |
| | Many cells/prominent ballooning | 2 |
| Fibrosis | None | 0 |
| | Perisinusoidal or periportal | 1 |
| | Perisinusoidal & portal/periportal | 2 |
| | Bridging fibrosis | 3 |
| | Cirrhosis | 4 |

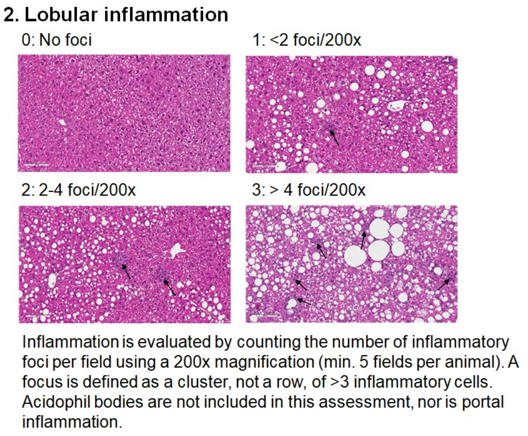


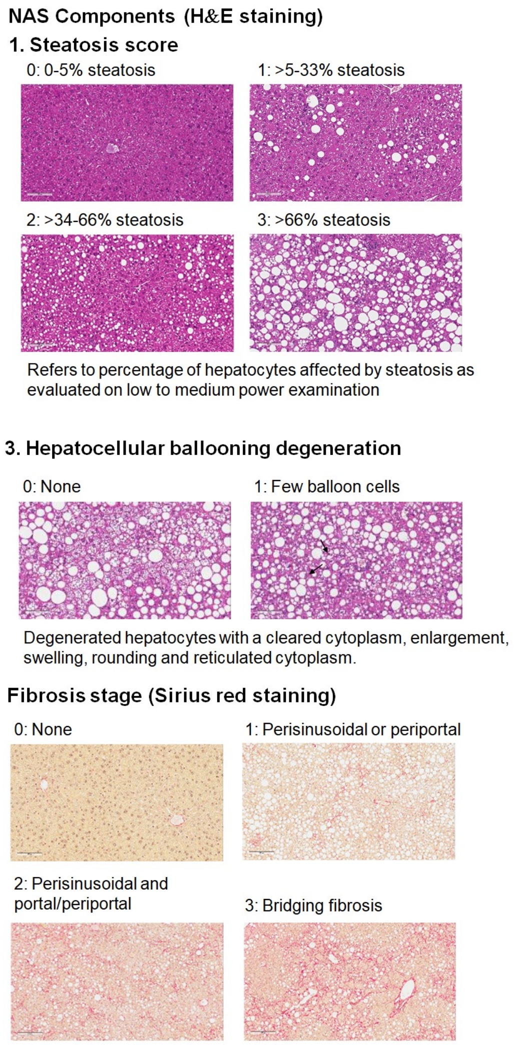


**Supplementary Fig. 3** Evaluation of fibrosis stage using Sirius Red staining and non-alcoholic fatty liver disease activity score (NAS) in hepatic sections stained with hematoxylin and eosin. The scores for steatosis, lobular inflammation, and hepatocellular ballooning degeneration are shown.

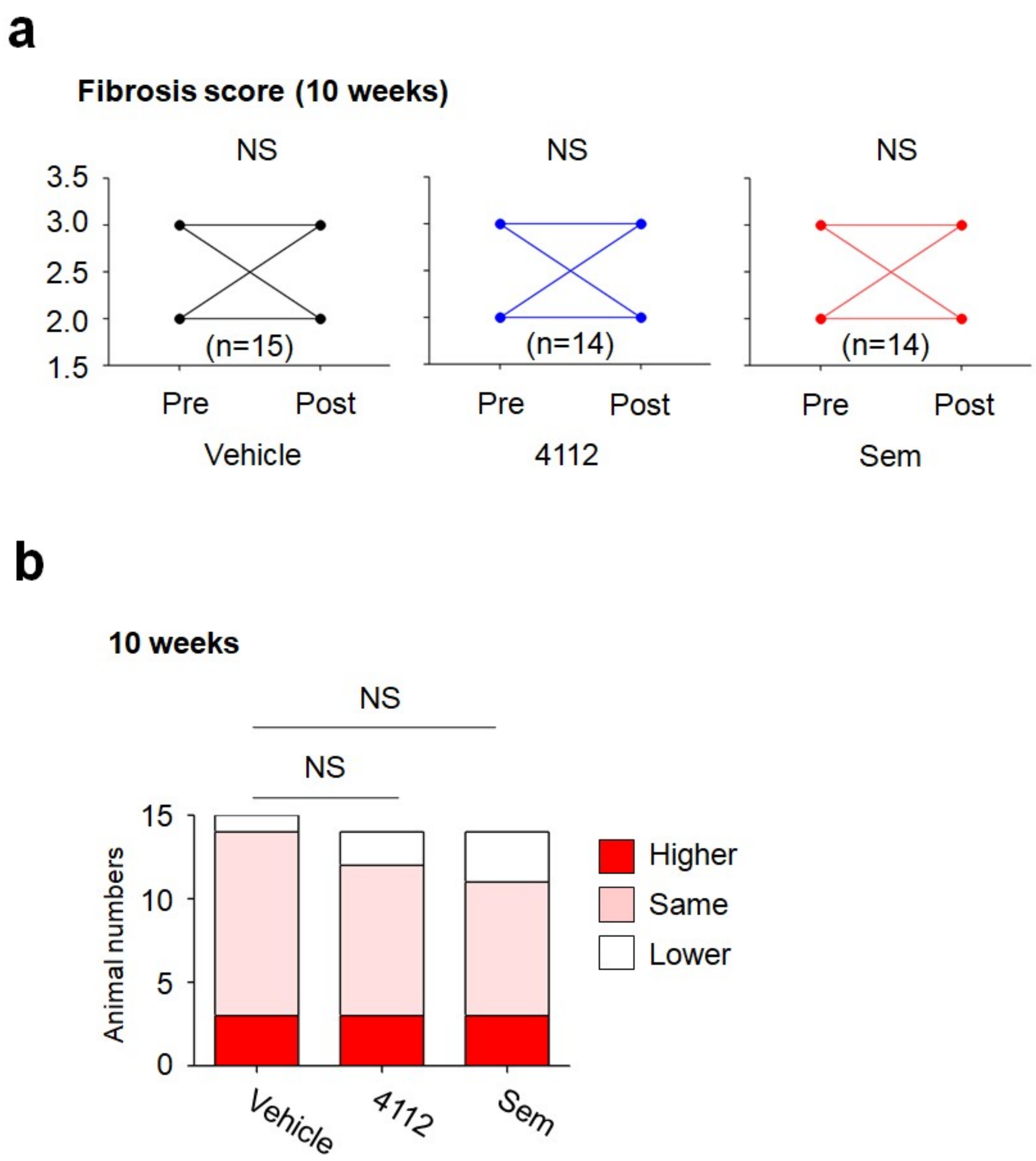


**Supplementary Fig. 4** Scoring of fibrosis stage after treating the biopsy-confirmed non-alcoholic steatosis hepatitis mouse model with drugs for 10 weeks (long-term). **a** Histological scores of fibrosis in biopsies before treatment (pre) and at the end of drug treatment (post). **b** The number of animals exhibiting exacerbation (), no change (same), and alleviation (lower) of disease after treatment relative to pre-treatment.

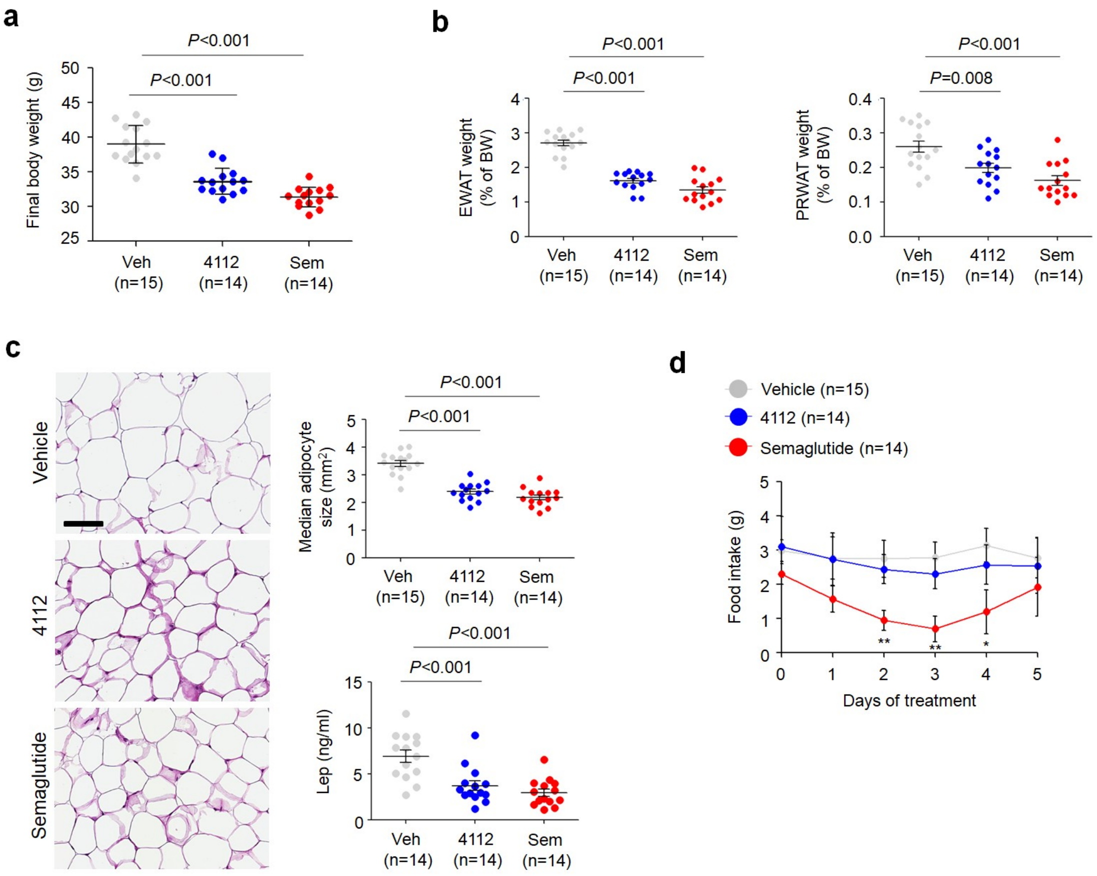


**Supplementary Fig. 5** Effect of HSG4112 and semaglutide on the final bodyweight (a) and the weights of epididymal white adipose tissue (EWAT) and pre-renal white adipose tissue (PRWAT) (b). **c** Histological analysis of white adipose tissues. Hematoxylin and eosin-stained adipocytes from white adipose tissue are shown on the left. Quantification of adipocyte size and leptin expression are shown on the right. **d** Daily food intake of HSG4112-treated and semaglutide-treated mice.

| | Sem/Veh | | 4112/Veh | |
|---|---|---|---|---|
| Lipid Class | FC | *P*-value | FC | *P*-value |
| MG | 0.61 | 0.00 | 0.78 | 0.00 |
| DG | 0.66 | 0.00 | 1.00 | 0.98 |
| TG | 0.26 | 0.00 | 0.73 | 0.04 |
| CE | 0.71 | 0.01 | 0.50 | 0.00 |
| Cholesterol | 0.74 | 0.00 | 0.81 | 0.03 |
| Cer | 0.90 | 0.20 | 0.97 | 0.87 |
| SM | 1.01 | 0.94 | 1.20 | 0.03 |
| Sa | 0.89 | 0.34 | 0.96 | 0.72 |
| So | 0.98 | 0.83 | 0.95 | 0.36 |
| PC | 1.05 | 0.22 | 1.07 | 0.10 |
| **LPC** | **1.20** | **0.59** | **0.86** | **0.01** |
| PE | 1.14 | 0.37 | 1.04 | 0.74 |
| **LPE** | **2.35** | **0.37** | **0.73** | **0.08** |
| PA | 1.02 | 0.66 | 1.08 | 0.13 |
| PI | 1.12 | 0.13 | 1.20 | 0.09 |
| **LPI** | **2.91** | **0.33** | **0.88** | **0.17** |
| PG | 1.01 | 0.95 | 1.06 | 0.59 |
| **LPG** | **1.60** | **0.46** | **0.91** | **0.39** |

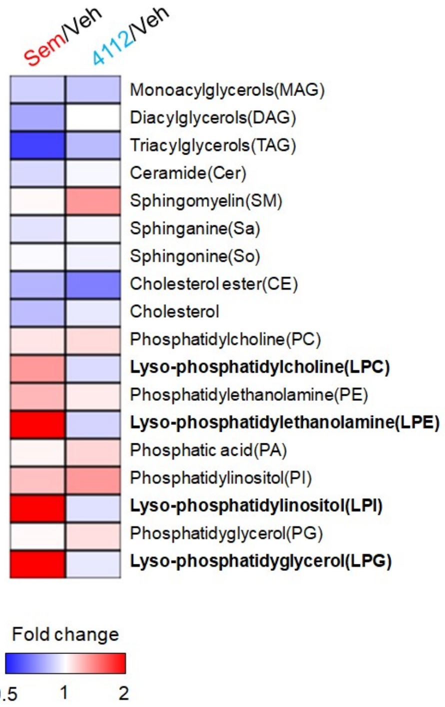


**Supplementary Fig. 6** Summary of lipid class analyses in the semaglutide-treated and HSG4112-treated amylin diet-induced non-alcoholic steatosis hepatitis model. Analysis of lipid class contents in the livers of HSG4112-treated and semaglutide-treated amylin fed mice is shown in the table and the heatmap. Each cell of the heatmap is color-coded based on the fold change of lipid levels in the drug-treated group relative to the vehicle-treated group. The legend for the color-coding is shown below.

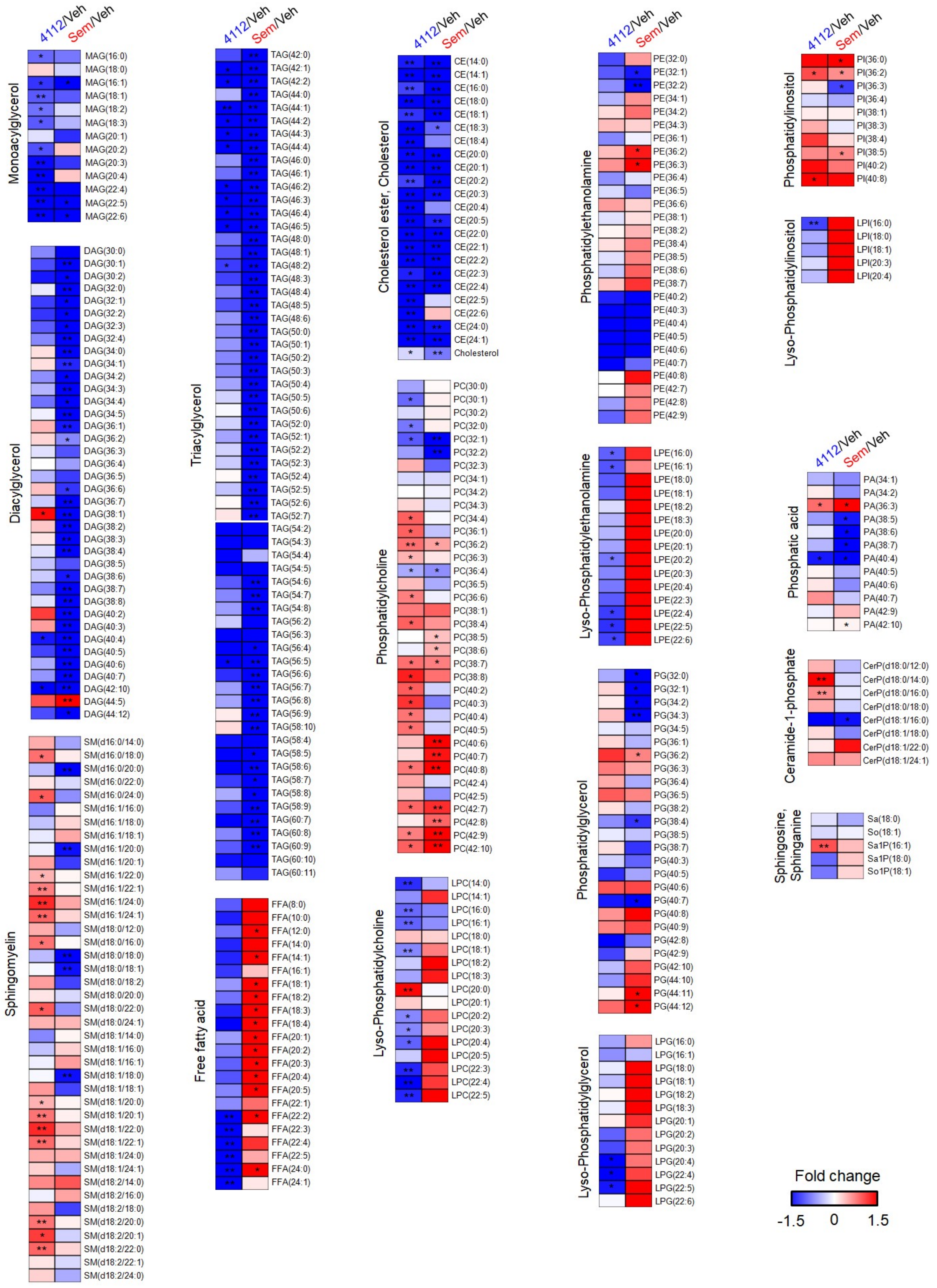


**Supplementary Fig. 7** Changes in the levels of individual lipid species in the semaglutide-treated and HSG4112-treated amylin diet-induced non-alcoholic steatosis hepatitis model. Heatmap analysis of individual lipid species in the liver of the HSG4112-treated and semaglutide-treated groups. Each cell is color-coded based on the fold change in the levels of individual lipid species in the drug-treated group relative to the vehicle-treated group. The legend for the color-coding is shown below. * $p < 0.05$, ** $p < 0.01$; drug-treated group vs. vehicle-treated group.

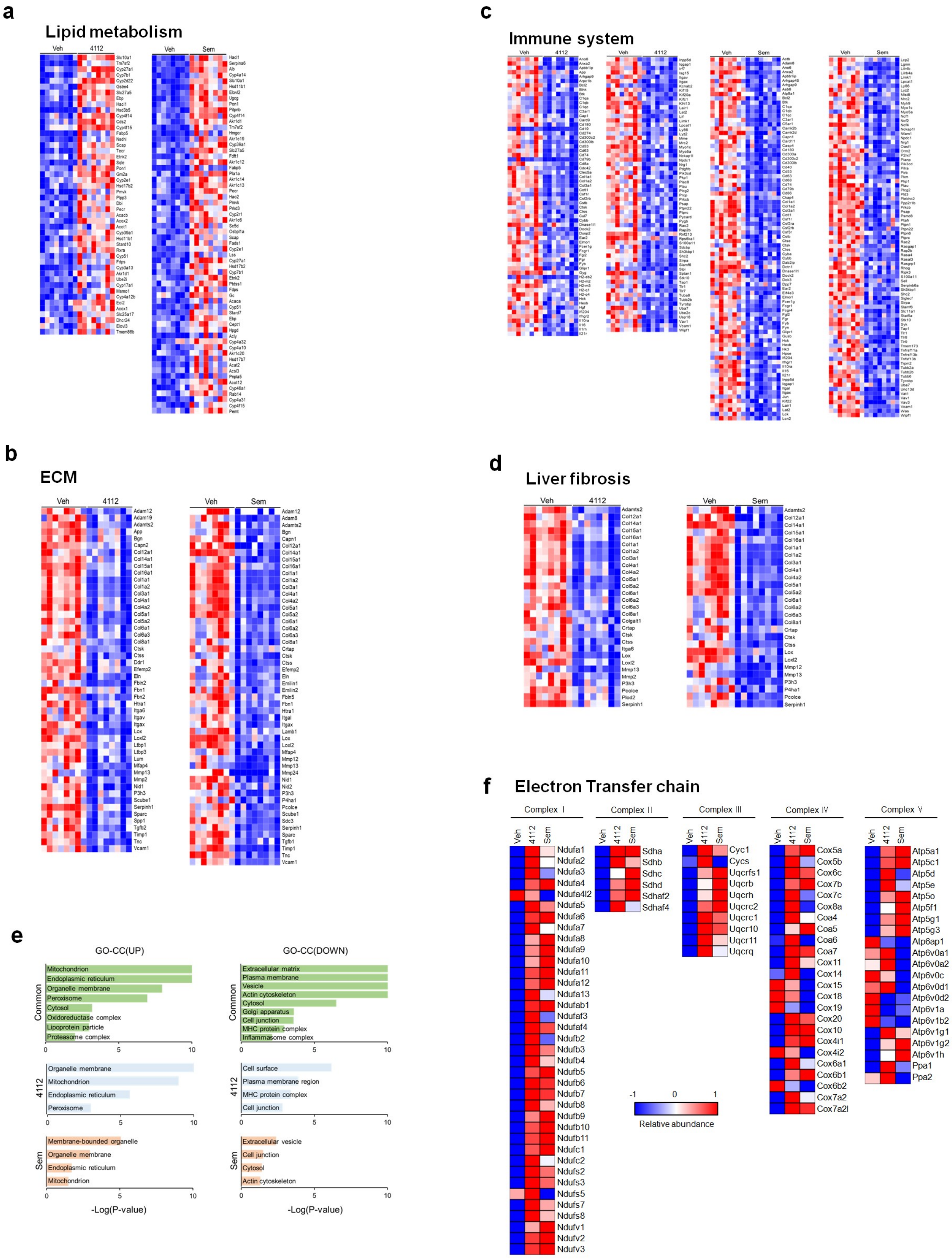


**Supplementary Fig. 8** Cellular component and heatmap analyses of RNA sequencing data of the HSG4112-treated and semaglutide-treated groups. **a** Heatmap showing the differentially expressed genes (DEGs) associated with lipid metabolism in the liver of

HSG4112-treated and semaglutide-treated amylin diet fed mice relative to the vehicle-treated group. Genes upregulated or downregulated by more than 2-fold are shown in red and blue, respectively. **b–d** Heatmap showing the DEGs associated with extracellular matrix (ECM) (b), immune system (c), and liver fibrosis (d). **e** Gene Ontology (GO) cellular component enrichment analysis. The GO cellular components of upregulated and downregulated genes in the liver of the HSG4112-treated and semaglutide-treated mice relative to the vehicle-treated group. **f** Heatmap showing DEGs involved in electron transfer chain in the HSG4112-treated and semaglutide-treated mice relative to the vehicle-treated group. Genes upregulated or downregulated by more than 2-fold are shown in red and blue, respectively. Each cell is color-coded based on the relative abundance of genes in the drug-treated group relative to the vehicle-treated group. The legend for the color-coding is shown below.

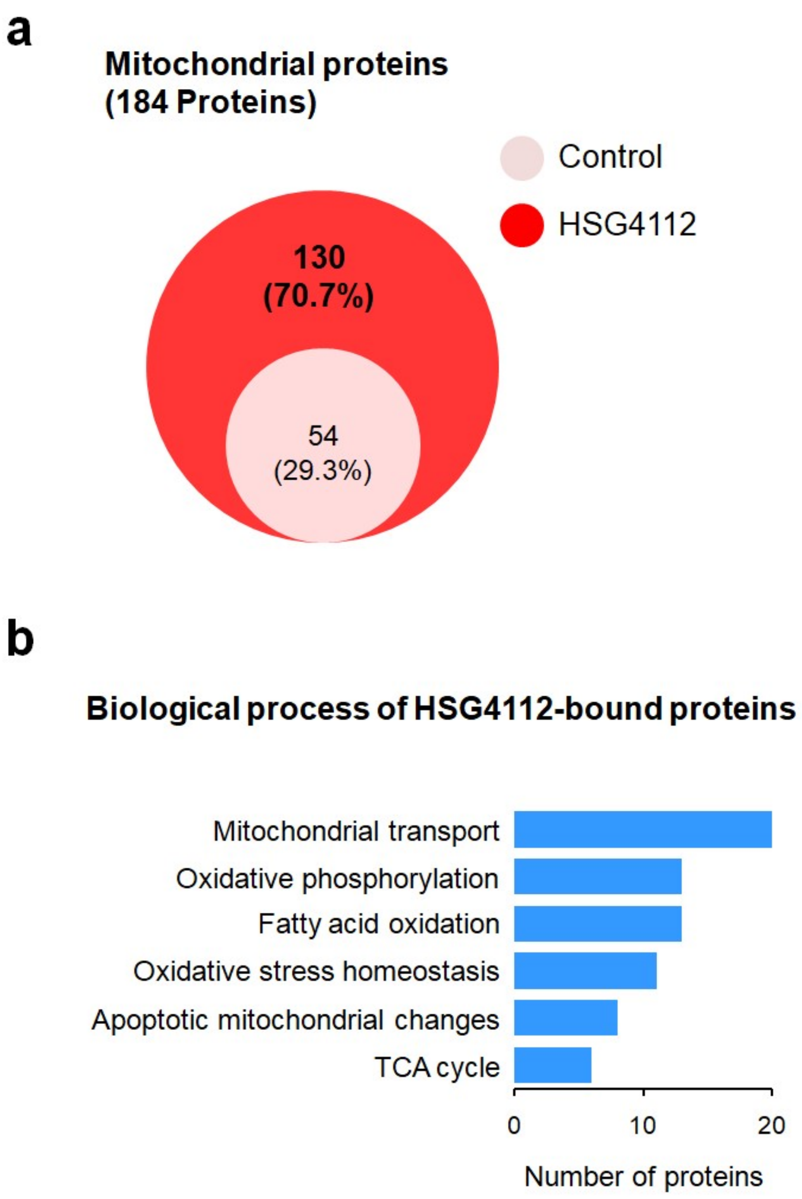


**Supplementary Fig. 9** Biological process analyses of HSG4112-bound proteins in the mitochondria. **a** Venn diagram showing the number of proteins bound with HSG4112-conjugated and control beads. **b** Gene Ontology (GO) biological process enrichment analysis. GO biological processes of HSG4112-bound proteins are related to the regulation of overall mitochondrial energy metabolism.
